# Multiomic State-Transitions Reveal Post-Treatment Transcriptome Desynchronization in Acute Myeloid Leukemia

**DOI:** 10.64898/2026.05.20.726707

**Authors:** Jennifer Rangel Ambriz, Ziang Chen, Yu-Hsuan Fu, David E. Frankhouser, Denis O’Meally, Lisa Uechi, Lianjun Zhang, Ying-Chieh Chen, Sergio Branciamore, Jihyun Irizarry, Bin Zhang, Guido Marcucci, Russell C. Rockne, Ya-Huei Kuo

## Abstract

Temporal dynamics of the peripheral blood transcriptome are crucial for understanding leukemia evolution and response to therapy because they can reveal how gene expression programs drive abnormal cell states, disease heterogeneity, and treatment resistance. Using a mathematical model of state-transitions, we studied the temporal dynamics of peripheral blood messenger RNA (mRNA) and microRNA (miRNA) transcriptomes in a mouse model of acute myeloid leukemia (AML). In our state-transition model, mRNA and miRNA transcriptomes are represented as a particle undergoing Brownian motion in a two-dimensional multiomic potential landscape. Following chemotherapy, we observed a temporal desynchronization between mRNA and miRNA transcriptomic responses corresponding to an asymmetric shift in the landscape. Specifically, mRNA trajectories responded almost immediately post-treatment, whereas miRNA responses were delayed by approximately two weeks. Clustering analysis identified that the temporal delay is driven by a prominent cluster of miRNAs from the imprinted *Dlk1-Dio3* region. Although previously implicated in acute promyelocytic leukemia, lymphomas, and metabolic dysregulation, this provides the first evidence linking the *Dlk1-Dio3* locus to AML chemotherapy response and treatment-induced transcriptomic desynchronization. This framework offers an innovative dynamics-based strategy to identify biological drivers of therapeutic response and novel therapeutic targets across hematological malignancies.

**Key Points:**

- Treatment can induce desynchronization between messenger RNA and microRNA transcriptome dynamics in a murine model of AML
- MicroRNAs from the imprinted *Dlk1-Dio3* locus are identified as key contributors to desynchronization and may serve as therapeutic targets

## Introduction

Biomarkers such as gene mutations and chromosomal abnormalities can be utilized for prognostic stratification and tailored treatment selection in acute myeloid leukemia (AML), a highly aggressive hematologic malignancy with long-term overall survival of approximately 30%^1^. However, current approaches that allow assessment of response to treatments require time, often revealing treatment ineffectiveness only after the patient has already shown treatment refractoriness or relapse. This creates a practical limitation in the clinic, as the window of opportunity to modify therapy early for non-responders may already be lost. Therefore, there is an unmet need for a systematic framework that can characterize disease dynamics and treatment response after initiation of a therapeutic intervention.

State-transition modeling is a useful framework to represent how a dynamic system moves between different states over time. Such models have been proven effective for understanding and predicting time-dependent behavior in stochastic systems and have been applied in a wide range of biological contexts including cellular differentiation, gene regulatory networks and disease^2–6^. We have previously shown that AML initiation and progression can be predicted with an application of state-transition theory to model movement of the transcriptome, both on the level of messenger RNA (mRNA) and microRNA (miRNA), in its corresponding state-space^7–10^. Leveraging time-series mRNA and miRNA sequencing data from peripheral blood (PB) collected during disease development, we constructed health-to-leukemia mRNA- and miRNA-based state-spaces and represented the dynamics of each transcriptome as a particle undergoing Brownian motion in a double-well potential energy landscape, characterized by mathematically defined critical points representing states of health and leukemia separated by a transition state^7–10^. We have also demonstrated the feasibility of applying a similar approach to assess and predict response of chronic myeloid leukemia (CML) to tyrosine kinase inhibitors (TKI)^9^, supporting the idea that transcriptomic state-transition represents a generalizable framework.

Here, we apply state-transition theory to characterize AML response to chemotherapy. We modeled mRNA and miRNA sequencing data obtained from time-sequential PBMC samples collected weekly from a murine model of inv(16) AML treated with chemotherapy. Herein, we report on a detailed longitudinal characterization of dynamic patterns of mRNA and miRNA during and after chemotherapy administration, guided by a state-transition mathematical model. This approach enables us to determine how transcriptomic state-transition track treatment response and disease relapse, while providing biological insights into mechanisms associated with treatment-induced transcriptome dynamics and disease outcome.

## Methods

### Mouse model

Conditional *Cbfb::MYH11* (CM) knock-in mice (*Cbfb*^*+/56M*^*/Mx1-Cre*; C57BL/6) were used as a model to recapitulate human inv(16) AML^11^, as previously reported^11,12^. All mice were maintained in an Association for Assessment and Accreditation of Laboratory Animal Care accredited animal facility, and all experimental procedures were performed in accordance with federal and state government guidelines and established institutional guidelines and protocols approved by the Institutional Animal Care and Use Committee (IACUC) at City of Hope.

### Experimental design for chemotherapy treatment and blood sampling

After detection of overt leukemia, which is monitored by the percentage of circulating leukemia blasts (cKit^+^>20%) detected by flow cytometry, CM mice (n=7) were treated with a “5+3” combination of cytarabine (Ara-C; 50mg/kg/day; 5 days) and daunorubicin (DNR; 1.5mg/kg/day; 3 days) to model the standard-of-care “7+3” chemotherapy for newly diagnosed AML patients^13^

(Figure 1). Similarly treated wild-type C57BL/6 mice (n=3) served as controls. PB was collected weekly from mice by retro-orbital bleeding^7^ before and after chemotherapy. The first post-treatment PB collection was two weeks after the initiation of treatment, as mice suffered from cytopenia and could not tolerate earlier sampling. Following red blood cell lysis, samples were subjected to flow cytometry and RNA extraction for bulk mRNA-sequencing (mRNA-seq) and miRNA-sequencing (miRNA-seq).

**Figure 1.**
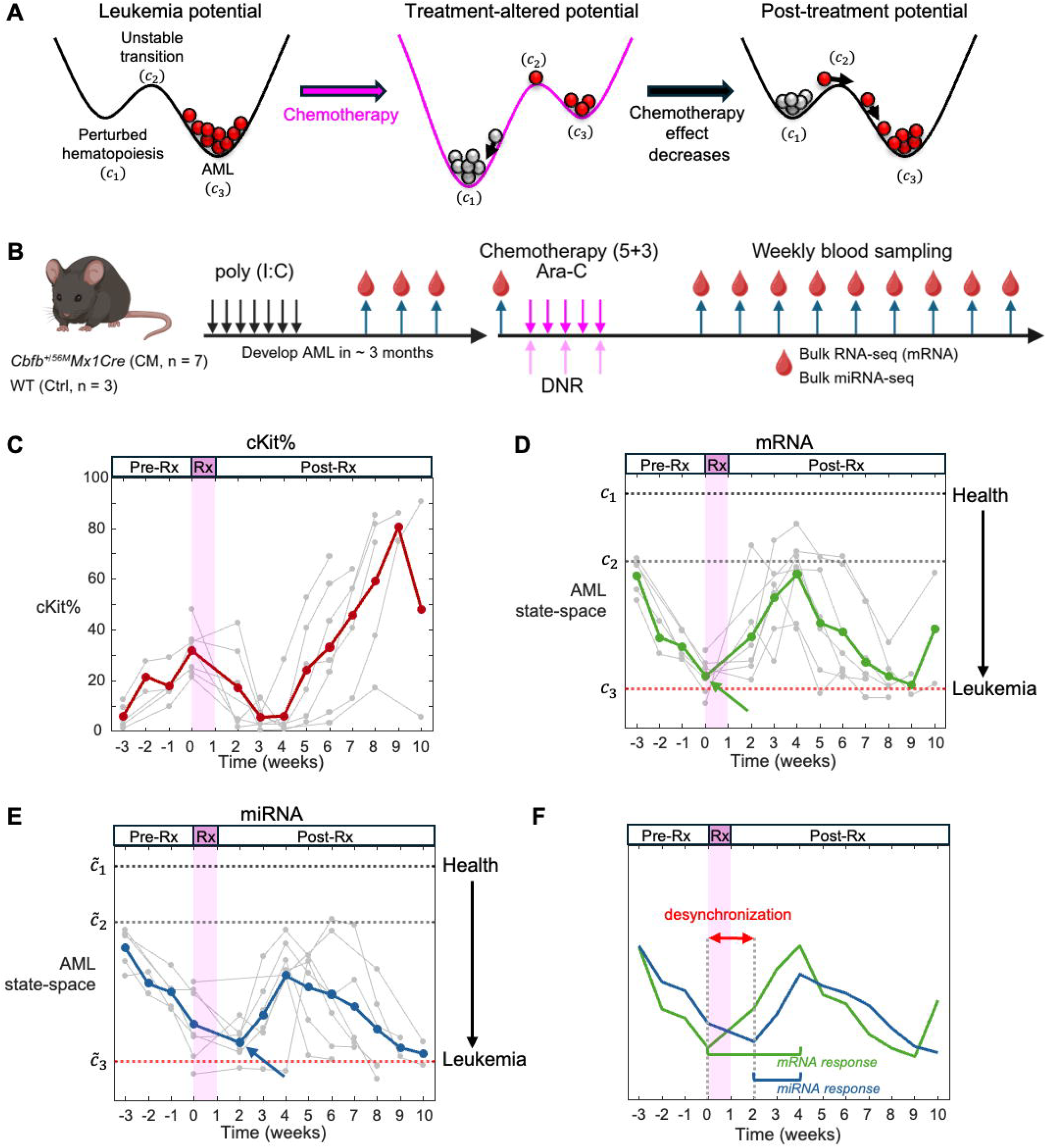
Longitudinal multiomic state-space trajectories reveal a temporal desynchronization of mRNA and miRNA transcriptome dynamics post-chemotherapy. **A)** A state-transition framework is applied to model each transcriptome (mRNA or miRNA) as a particle undergoing Brownian motion in a potential energy landscape defined by three critical points corresponding to health (*c*_1_), transition (*c*_2_), and leukemic (*c*_3_) states. **B)** Schematic of experiment design. Expression of *Cbfb::MYH11* (CM) in conditional CM knock-in mice (*Cbfb*^*+/56M*^*/Mx1-Cre*; C57BL/6) was induced with intraperitoneal injections of poly (I:C). Time-series PBMC samples from CM leukemic mice (n = 7) were collected weekly before and after a “5+3” chemotherapy regimen consisting of cytarabine (Ara-C) and daunorubicin (DNR). Samples were analyzed by bulk RNA-seq and miRNA-seq. Created in BioRender Rangel Ambriz, J. (2026) https://BioRender.com/bwr8f3n. **C)** Temporal dynamics of circulating cKit+ cells were monitored weekly. Chemotherapy was initiated when circulating leukemia blasts exceeded 20% (cKit^+^ > 20%), indicating overt leukemia. The mean cKit% trajectory (red) shows an initial reduction of leukemic blasts post-treatment followed by subsequent rebound. **(D-E)** Longitudinal mRNA and miRNA transcriptome trajectories of seven treated CM mice (gray) projected into their respective reference AML state-spaces, which was constructed using untreated CM mice (2018 cohort). Chemotherapy treatment (pink) was initiated at W0 (Time 0). Mean mRNA (green) and miRNA (blue) trajectories summarize the transcriptome dynamics relative to state-transition critical points, showing movement from the leukemia state (*C*_3_ and 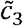) towards a healthy state (*C*_1_ and 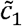) post-chemotherapy, followed by relapse toward the leukemic state (*C*_3_ and 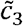). The start of treatment response of each trajectory is denoted with an arrow. **F)** Overlay of mean trajectories reveals a temporal desynchronization of the mRNA and miRNA transcriptome dynamics post-chemotherapy, where the mRNA (green) trajectory shows an immediate response to chemotherapy meanwhile the miRNA trajectory (blue) exhibits a delayed response.

### RNA sequencing and data processing

PB RNA was extracted, inspected for quality, and used for mRNA-seq and miRNA-seq library preparation, followed by high-throughput sequencing on Illumina platforms. Reads were processed using nf-core RNASeq (v3.19.0) and smRNAseq (v2.3.1)^14^, aligned to the GENCODE primary mouse reference (release M33) amended with human transgenes to generate gene- and miRNA-level expression matrices and quality assessment (Supplemental Information).

### State-space projection of chemotherapy-treated mice

We used principal component analysis (PCA) to project the transcriptome states of chemotherapy-treated mice into previously defined mRNA- or miRNA-based reference state-spaces from untreated CM and control mice^7,8^. Briefly, reference state-spaces were constructed by performing PCA via singular value decomposition on mean-centered, log-transformed mRNA or miRNA expression matrices. As previously reported, PC1 and PC2 define the miRNA- and mRNA-based reference state-spaces, respectively, since these PCs most strongly correlated with *Kit* expression and resulted in the largest separation between CM and controls^7,8^. Loading values of each expression matrix were used to project the treated CM data into the reference AML state-spaces (Supplemental Information).

### 2-Dimensional (2D) state-transition model

We modeled mRNA-miRNA state-transition dynamics as a particle undergoing Brownian motion in a 2D potential energy landscape with a Langevin equation of the form:

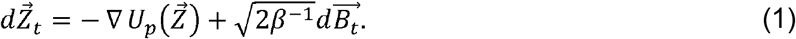

Position of the particle in the potential energy landscape 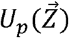 is defined by the vector 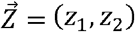, where z_1_ and z_2_ represent the position of the mRNA and miRNA transcriptomes in the state-space, respectively. *B*_t_ is a Brownian process, with diffusion coefficient β, estimated by mean-squared displacement analysis of state-space trajectories (Figure S1).

In untreated disease progression, the leukemic potential landscape is defined as

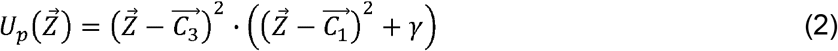

where 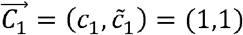 and 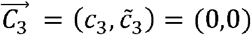 denote the health and disease states, respectively. Critical points *c*_1_ and *c*_3_ corresponding to the mRNA-based state-space, and 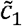 and 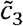, corresponding to the miRNA-based state-space, were estimated from untreated CM mice in prior work^7,8^. βrepresents the effect of the CM oncogene, altering the shape of the potential to give a higher probability of the disease state (Figures S2-3).

Chemotherapy effects were modeled as time-dependent deformations of the 2D potential landscape, represented mathematically as:

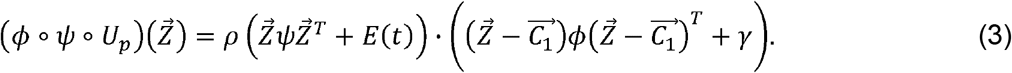

Here, ψ and ϕ are 2×2 tensor operators that map the effects of disease and chemotherapy on the potential landscape, respectively, *T* is the transpose, ρ is a scaling coefficient and *E*(*t*) is the pharmacokinetic-pharmacodynamics (PK-PD) of chemotherapy (Supplemental Information).

### A state-space defined treatment vector

To quantify how chemotherapy perturbs the mRNA-miRNA dynamics and deforms the potential landscape, we defined a treatment vector 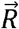 as the difference between untreated and treatment-perturbed gradients of the potential fields. The largest change in direction is given by position in the state-space that results in the maximum angular difference between two vector fields (Figure S4). This vector quantifies the direction and magnitude of chemotherapy-induced landscape deformation, providing a quantitative measure of how treatment affects the mRNA and miRNA transcriptomes (Supplemental Information).

### State-space based sample grouping and expression analysis

Because disease progression rates varied across mice, mRNA samples from each mouse at each timepoint were clustered into three groups based on their state-space proximity to critical points (Figure S5A): pre⍰treatment leukemic state near *c*_3_ (Pre⍰Rx), maximal treatment response near *c*_1_ (Recovery), and relapse back toward *c*_3_ (Relapse). To identify differentially expressed genes (DEGs) and miRNA among the three groups, DESeq2^15^ was applied with default parameters using gene count data generated via tximport^16^ from Salmon abundance estimates^17^. Subsequently, Gene Set Enrichment Analysis (GSEA)^18,19^ was performed utilizing the MSigDB hallmark databases^20^ to identify pathways perturbed before and after chemotherapy.

### Co-expression cluster analysis

Top 25% of mRNAs or miRNAs with highest variance among all samples were used for Weighted Gene Co-expression Network Analysis (WGCNA)^21^ to identify co-expressed mRNAs or miRNAs clusters. Identified clusters were projected back into the respective state-space to assess each cluster’s contribution to disease progression and chemotherapy response

(Supplemental Information). Location of miRNAs in the genome was visualized using PhenoGram^22^.

## Results

### Construction of state-spaces reveals desynchronization of mRNA and miRNA state-transition dynamics after chemotherapy

We applied a state-transition framework to investigate the dynamics of the mRNA and miRNA transcriptomes following chemotherapy treatment defined by three critical points that correspond to a health state (*c*_1_), a transition state (*c*_2_), and a leukemic state (*c*_3_) (Figure 1A)^7,8^.

We hypothesized that chemotherapy perturbs the potential landscape, reducing the energy barrier and thereby increasing the probability of a transition from a leukemic state back toward a health state. To test this hypothesis, we collected longitudinal PB samples from a conditional CM knock-in mouse model that recapitulates inv(16) AML^11,12^ treated with chemotherapy (Figure 1B). Chemotherapy initially reduced the number of cKit^+^ cells in the PB, but all mice eventually relapse (Figure 1C).

To conduct a state-transition analysis, we constructed mRNA and miRNA state-spaces in which we could observe time-series disease trajectories. We then plotted the mRNA and miRNA transcriptome state-space trajectories over time and used a mean trajectory to summarize the dynamics relative to state-transition critical points during disease progression and following therapeutic treatment (Figure 1D-E).

Prior to treatment, both mRNA and miRNA transcriptome trajectories moved synchronously from a health state (*c*_1_ and 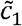) toward a leukemic state (*c*_3_ and 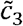). Following chemotherapy, the transcriptome trajectories moved away from leukemic state toward the healthy state before returning to the leukemic state upon disease relapse (Figure 1D-E). We observed that the mRNA and miRNA transcriptomes moved differently relative to the critical points in the state-space following chemotherapy. Notably, there was a delayed response of the miRNA transcriptome trajectories as compared with the mRNA transcriptome, which we hereafter refer to as a *desynchronization* between the mRNA and miRNA transcriptome dynamics (Figure 1F, Figure S6).

Specifically, the mRNA trajectories moved from *c*_3_ towards *c*_1_ immediately after treatment, whereas the miRNA transcriptome trajectories continued to progress towards 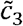 for 2 or more weeks after treatment, before recovering. The trajectories synchronized again approximately 4 weeks after treatment, coinciding with disease relapse, when both transcriptome dynamics aligned and progressively returned toward the leukemic state. Comparing the percentage of cKit+ leukemia cells with both transcriptome trajectories before and after chemotherapy, we observed that prior to treatment and during AML progression, both mRNA (*R*^2^= 0.72) and miRNA (*R*^2^ = 0.74) were strongly correlated with the percent of cKit+ cells in the PB as measured by flow cytometry (Figure S7). Following chemotherapy, the correlations decreased, with a substantially greater reduction for miRNA (*R*^2^ = 0.30) than for mRNA (*R*^2^= 0.67).

### Multiomic state-transition framework captures mRNA-miRNA dynamics post-chemotherapy

Since we identified that mRNA and miRNA transcriptome dynamics were desynchronized following chemotherapy, we extended our state-transition model to a 2D framework to simultaneously model mRNA-miRNA transcriptome dynamics and better understand how their dynamics evolve across different disease states (Figure 2A). To do so, we constructed a multiomic state-space by pairing the mRNA and miRNA transcriptome data from individual untreated and chemotherapy-treated CM mice at each time-point, enabling simultaneous assessment of disease state across both modalities.

**Figure 2.**
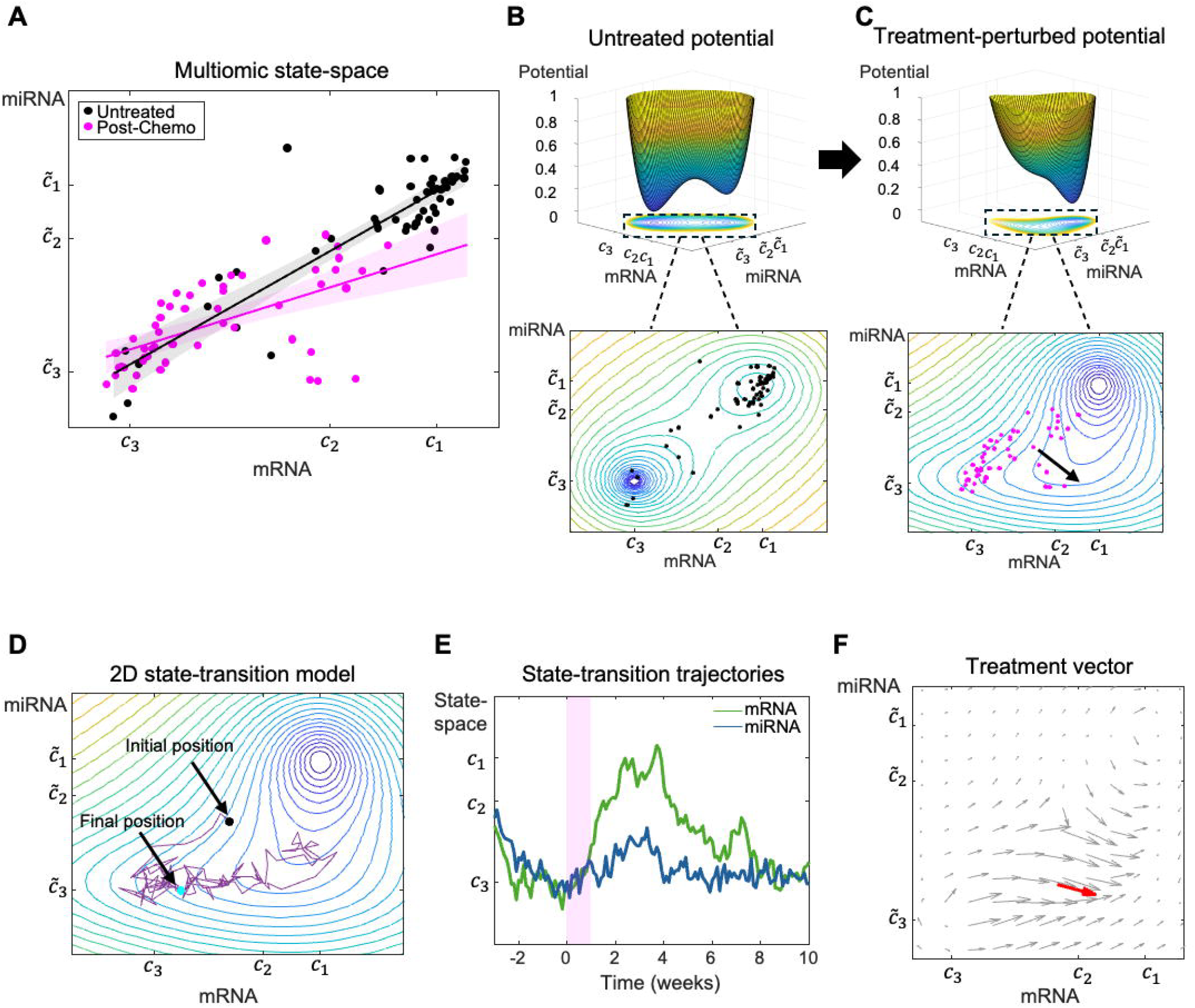
Multiomic state-space and 2D state-transition model. **A)**Multiomic state-space was constructed by pairing the mRNA and miRNA transcriptome data based on timepoint from individual untreated (black) and post-treatment CM mice (magenta). The post-treatment time-points include the start of chemotherapy up to 10 weeks post-treatment. There is a strong correlation (*R*^2^=0.77) between the mRNA-miRNA dynamics during disease progression and in the absence of treatment meanwhile, there is a poor correlation in chemotherapy-treated mice dynamics (*R*^2^ = 0.35). **B)** Top panel: Shape of leukemic potential landscape in three-dimensions (3D) under untreated conditions with contour plots shown underneath. The leukemic potential is characterized by two wells of unequal depth, each representing a state of health (*C*_1_, 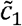)or disease (*C*_3_, 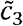). The most energetically favorable state (*C*_3_, 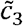) is the disease state, which has a deeper well to reflect the increased probability of a health-AML transition during disease progression and under no treatment conditions. Bottom panel: Contour plots of the untreated potential landscape capture the correlated mRNA-miRNA relationship from the untreated multiomic state-space data (black). **C)** Top panel: multiomic potential landscape during maximum chemotherapy effect with contour plot shown underneath. Chemotherapy acts as an anti-leukemic force that shifts the leukemic landscape to make the health state more energetically favorable by increasing the depth of the (*C*_1_, 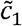 well and in turn, increasing the likelihood of a disease-health transition. Bottom panel: Contour plots of the potential landscape during maximum chemotherapy effects (*E=E*_*max*_) are shown overlaid with the multiomic state-space data. The deformation of the landscape is in the same direction (black arrow) as the chemotherapy-treated mice data (magenta). **D)** The stochastic equation of motion was solved forward in time for the leukemic and chemotherapy-treated potentials to predict the trajectory of a 2D particle with position 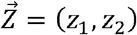 in the multiomic state-space (purple trajectory). The initial and final positions of the simulated mRNA-miRNA trajectory are denoted in black and cyan, respectively. The trajectory is overlaid on the simulated contour plots of the potential landscape during maximum chemotherapy effects. **E)** The mRNA (green) and miRNA (blue) trajectories over time generated by the 2D state-transition model. **F)** To quantify how chemotherapy affects the mRNA and miRNA transcriptomes, we quantified the effect of treatment by evaluating the difference between the untreated vector field and the treatment-perturbed vector field during maximum treatment effect resulting in a treatment vector 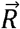(red vector). The difference field over the entire state-space is denoted with gray vectors.

To model how chemotherapy alters the leukemic potential landscape (Figure 2B), we incorporated a time-dependent chemotherapy effect *E*(*t*) derived from a PK-PD model^23,24^ that accounts for the drug dosage and plasma half-life concentrations of Ara-C and DNR reported in literature^25,26^ (Supplemental Information). We also introduced two 2×2 tensors, ψ and ϕ, that mathematically and geometrically encode how CM and chemotherapy deform the 2D landscape (Figure 2C).

With the potential landscape and model parameters estimated from the data, we simulated transcriptomic trajectories evolving under chemotherapy (Figure 2D, Video S1). These simulations allowed us to observe how mRNA-miRNA dynamics evolve over time as chemotherapy deforms the potential landscape. The resulting mRNA-miRNA trajectory demonstrates how treatment-induced desynchronization drives transitions between leukemic and healthy states. When we plotted single-modality trajectories generated by the 2D state-transition model, we found that simulated trajectories transition from leukemic to healthy states post-treatment but eventually relapse, recapitulating the temporal behavior observed in the experimental data (Figure 2E).

Finally, to quantify how chemotherapy affects the potential landscape we computed a treatment difference vector 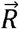 (Figure 2F). Notably, 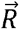 points in the southeast direction in the state-space, demonstrating that chemotherapy asymmetrically affects the mRNA transcriptome, consistent with the observed desynchronization.

### State-space analysis reveals dynamic immune and metabolic pathway reprogramming during treatment response and relapse

To characterize functional changes, we analyzed pathway dynamics across disease progression and treatment response relative to state-transition critical points: Pre-Rx, Recovery, and Relapse (Figure S5A). The relapse state showed the greatest number of DEGs as compared to treated controls (Figure S5B, Table S1). Pathway dysregulation, including proliferative (E2F targets, G2M checkpoint), estrogen response, and apical surface pathways, were observed even at maximal treatment response (Recovery vs Control; Figure S5C), indicating incomplete molecular restoration. Comparison between relapse and pre-Rx groups (Table S1), identified 1,571 dysregulated genes (Figure S5D), enriched in interferon signaling, inflammatory response, and TNFα/NF-κB pathways, indicating immune reprogramming specific to relapse post-chemotherapy (Figure S5E).

We examined changes in gene expression and pathway activity at each state using Gene Set Variation Analysis (GSVA) and identified dysregulation across several key biological programs (Figure S5F). This analysis revealed coordinated dysregulation of multiple biologically distinct programs. Immune and inflammatory signaling pathways, including TNFα signaling via NF-κB, IL-6–JAK–STAT3, and interferon response programs, were markedly upregulated post-chemotherapy and subsequently downregulated at relapse, suggesting a strong immune response to chemotherapy that gradually diminishes as the disease relapses. These pathways have been shown to promote AML cell growth and have been associated with chemoresistance^27–29^. Key metabolic pathways associated with AML progression^30–36^ like oxidative phosphorylation (OXPHOS), glycolysis and fatty acid metabolism were downregulated during the recovery stage post-chemotherapy, but were upregulated during relapse.

Hierarchical clustering of GSVA pathway scores identified three major dynamic groups that change together post-chemotherapy (Figure 3A). Three groups showed clear transitions following treatment: Group 1 had a local minimum at week 0, whereas Group 3 exhibited a pronounced peak at this time (Figure 3B). Groups 2 and 3 dropped markedly by week□ and were enriched for metabolic pathways commonly dysregulated in cancer and associated with AML progression and chemoresistance^30–36^, including OXPHOS, PI3K–AKT–mTOR signaling, and reactive oxygen species (ROS) pathways. Individual pathway dynamics also confirmed the suppression of these metabolic programs post-chemotherapy (Figure S8).

**Figure 3.**
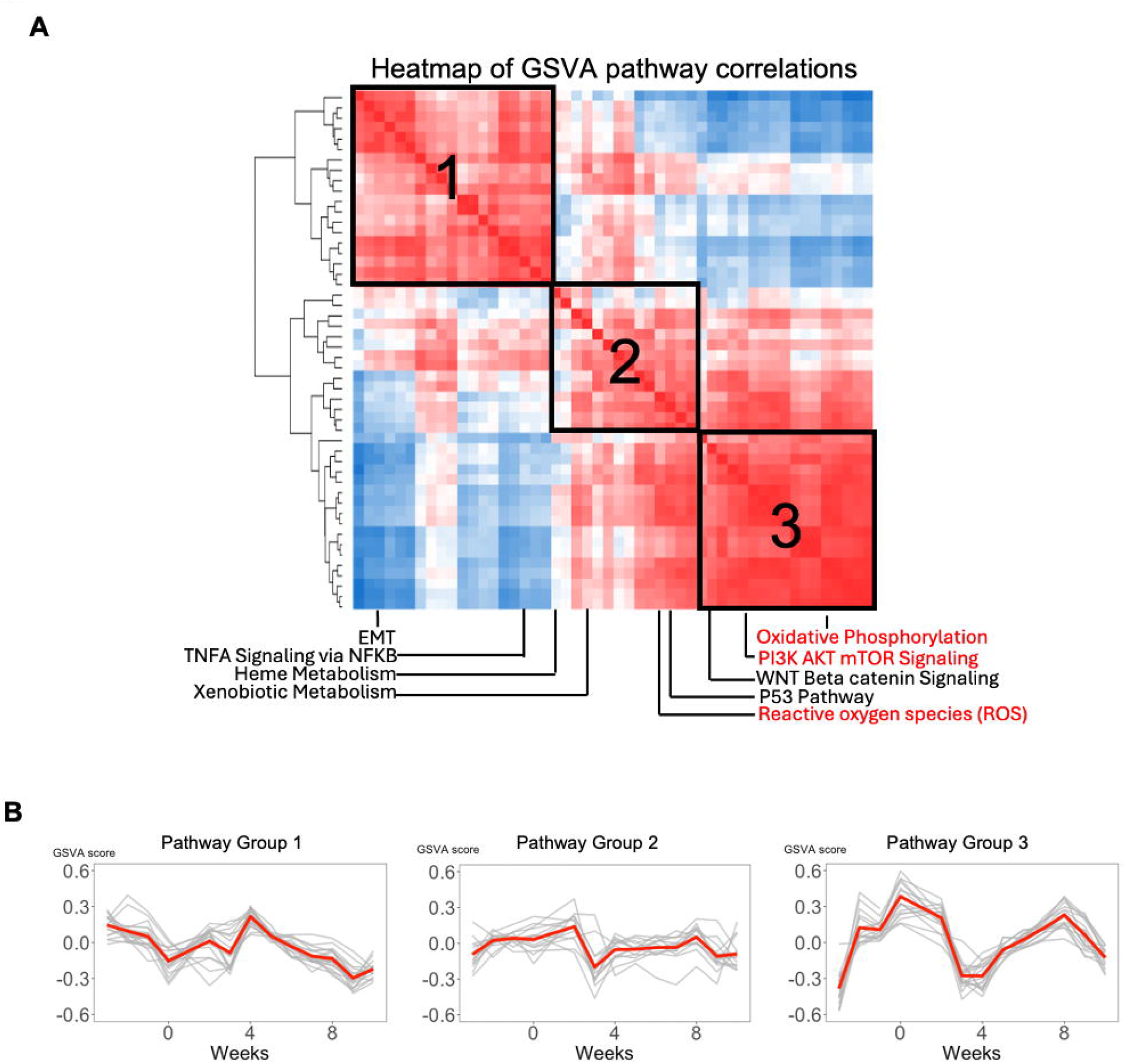
Temporal Dynamics of Pathway Activity Revealed by GSVA Highlight Distinct Clusters and Metabolic Suppression. **A)** Hierarchical clustering of pathway activities scored by GSVA across time. Four distinct dynamic pathway patterns were identified. Metabolism pathways are marked in red (Figure S8). **B)** Mean expression trajectories of each group pathway dynamics (red). Gray lines represent a pathway’s temporal GSVA score over time.

### Co-expression network analysis reveals distinct but coordinated mRNA and miRNA dynamics

To gain biological insight into the delayed miRNA transcriptome response relative to mRNA following chemotherapy, we applied WGCNA on both expression profiles and projected the resulting co-expression modules onto the leukemia state-space (Figure 4; Tables S2-3). WGCNA of mRNA expression profiles identified nine co⍰expression clusters (labeled a-i) based on their temporal expression dynamics. To assess each cluster’s contribution to leukemia progression and treatment response, we analyzed PCA loading values of the mRNA expression matrix (Figure 4A), where negative loading values indicate pro-leukemic effects and positive values indicate anti-leukemic effects. Most of these clusters showed negative loading values, contributing to overall pro⍰leukemic activity. In parallel, WGCNA of miRNA expression profiles identified seven clusters (labeled 1-7), of which only two clusters (clusters 1,2) exhibited strong pro-leukemic loading values (Figure 4B).

**Figure 4.**
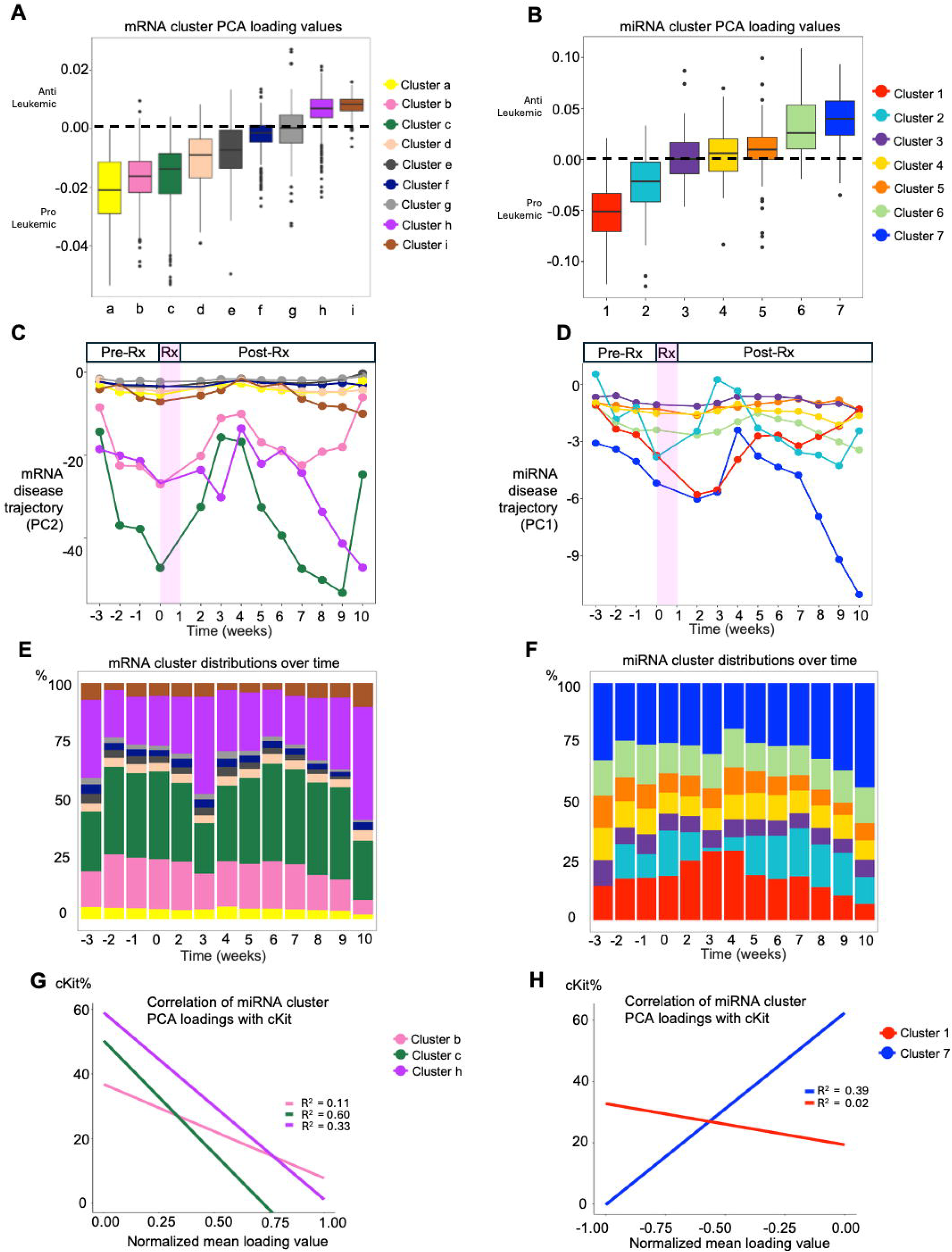
Weighted gene co-expression network analysis identifies miRNA and mRNA clusters with distinct temporal dynamics. **A)**Weighted Gene Co-expression Network Analysis (WGCNA) identified 9 mRNA clusters based on similar temporal expression dynamics. Average PCA loading values in each cluster were computed to assess the pro- or anti-leukemic effects of the expression on state-transition. A negative loading value indicates that increased expression of the corresponding miRNA results in a trajectory toward *c*_3_ in the state-space and is therefore pro-leukemic and vice versa for anti-leukemic. **B)** The same WGCNA analysis for miRNAs identified 7 miRNA clusters. **C)** Trajectories of each mRNA cluster were obtained by projecting each cluster into the untreated mRNA state-space. **D)** Trajectories of miRNA clusters using the same analysis as in C. **E)** The contribution of each mRNA cluster to the location of the samples in the state-space is computed as the fraction of the total state-space locations per timepoint. **F)** The contribution of each miRNA cluster. **G)** Correlation between normalized mean PCA loading value with cKit+%. Strong correlations with cKit+% (*R*^2^ > 0.5) suggest the mRNAs within the specified cluster have a strong association with AML disease dynamics. Correlation plots with data points for individual clusters can be found in Figure S13A **H)** Correlation between cKit+% and selected miRNA clusters. Correlation plots with data points for individual clusters can be found in Figure S13B

Projection of individual clusters onto the mRNA state-space identified clusters b (394 mRNAs), c (660 mRNAs), and h (2,293 mRNAs) as the greatest contributors of bulk mRNA dynamics (Figure 4C-E). Clusters b and c exhibited an immediate transcriptional response following chemotherapy. Both clusters had negative average loading values, indicating that increased expression of these clusters drives movement toward the leukemic state. Since the expression of these mRNAs decreased sharply after treatment, the overall mRNA trajectory shifts immediately toward the healthy state. Pathway enrichment analysis showed that these clusters were enriched for cell proliferation, mitotic, and hormonal signaling programs (Figure S9). In contrast, mRNA cluster h displayed a delayed response, with transient downward movement toward the leukemic state followed by recovery. Cluster h was enriched for immune and inflammatory pathways (Figure S9).

Projection of each miRNA cluster onto the miRNA state-space revealed that clusters 1 (30 miRNAs) and 7 (93 miRNAs) together dominate the miRNA trajectory 2-4 weeks post-chemotherapy, driving movement toward the leukemic state during this interval (Figure 4D-F). Computationally setting the expression of these two clusters to their pre-chemotherapy levels eliminated the delayed miRNA transcriptome response post-treatment (Figure S10), also suggesting that clusters 1 and 7 are the principal contributors to this delayed chemotherapy response. These clusters exhibited opposing functional roles, with miRNA cluster 1 (pro-leukemic) being upregulated, and miRNA cluster 7 (anti-leukemic) being downregulated (Figure S11). We next identified mRNA targets of miRNAs in each cluster using miRecords^37^, miRTarBase^38^, and TarBase^39^ databases. Pathway enrichment analysis of each of these targets revealed that cluster 1 targets were enriched for immune and stress-response programs, including TNFα/NF-κB, IL-2–STAT5, hypoxia, and TGF-β signaling, while cluster 7 targets were enriched for metabolic programs such as fatty acid metabolism, adipogenesis, and xenobiotic metabolism, implicating these miRNA clusters in microenvironmental signaling and metabolic adaptation, respectively (Figure S12).

Finally, correlating mRNA and miRNA clusters with cKit^+^ percentage revealed that mRNA clusters c and h, and miRNA cluster 7, correlated relatively higher with disease progression than mRNA cluster b and miRNA cluster 1 (Figure 4F-G). Together, these findings suggested that the transcriptomic state-space trajectory comprise dynamic clusters with heterogeneous relationships to disease progression and responsiveness to treatment (Figure S13), contributing to a desynchronization between mRNA and miRNA transcriptional trajectories during chemotherapy.

### Imprinted *Dlk1-Dio3* locus is associated with desynchronization of miRNA and mRNA transcriptomes

We next examined the distribution of co-expression clusters across chromosomes to identify regions that may be responsible for the observed desynchronization (Figure 5A-B). mRNA clusters were broadly dispersed across the genome, reflecting wider genomic distribution of protein-coding genes. While most miRNA clusters were broadly distributed across chromosomes, clusters 1 and 7 were each predominantly concentrated on a single chromosome.

**Figure 5.**
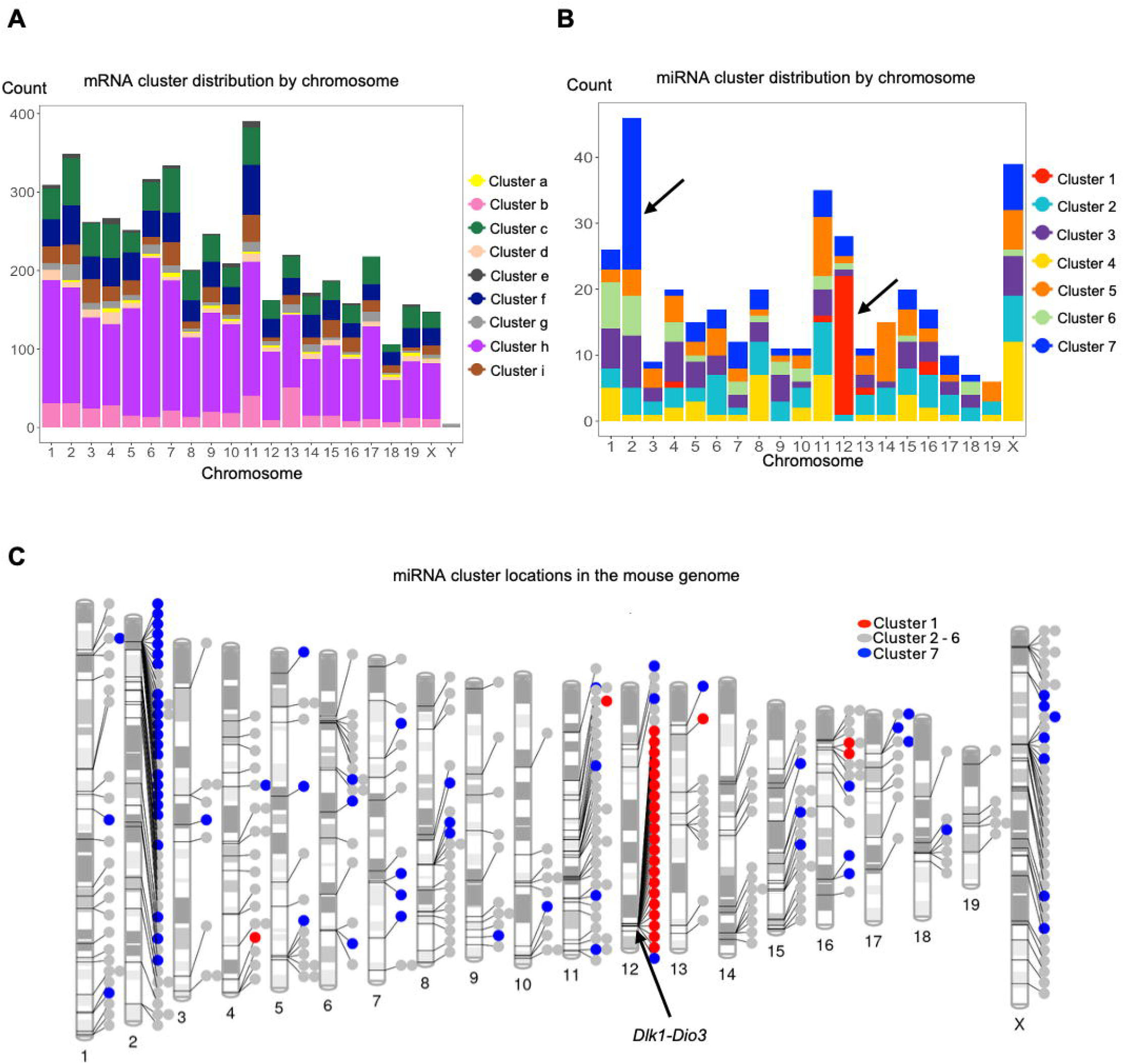
Genomic localization of miRNA and mRNA clusters and pathway dynamics across leukemia state transitions. Chromosomal distribution of **A)** mRNA and **B)** miRNA clusters. **C)** Genomic mapping phenogram of miRNAs by cluster across the mouse genome reveals that approximately 80% of the miRNAs in Cluster 1 are localized within the *Dlk1-Dio3* imprinted locus on chromosome 12 in mice (12qF1).

Cluster 1 miRNAs were predominantly localized to chromosome 12 with 25 of 30 miRNAs (80%) originating from this locus (Figure 5C). Notably, these miRNAs mapped into the *Dlk1-Dio3* imprinted locus at chromosome 12qF1 in mice (Table S4), representing a significant association between this locus and cluster 1 (p < 0.001). The *Dlk1-Dio3* locus is evolutionarily conserved in humans (14q32) and is known to produce coordinately expressed miRNAs under a shared regulatory elements^40^. miRNAs from the *Dlk1-Dio3* locus have previously been implicated in multiple disease context, including acute promyelocytic leukemia, lymphomas, glioblastoma, stem cell biology, stress response, and type 2 diabetes^40–42^. Consistent with prior reports linking this miRNA locus to the coordinately regulation of mitochondrial metabolism^33–35^, pathway analysis revealed broad downregulation of mitochondrial metabolic pathways post-chemotherapy (Figure S8).

miRNAs in cluster 7 predominantly map to a rodent-specific paternally imprinted genomic locus on chromosome 2 (p<0.001), corresponding to the intronic *Sfmbt2* miRNA cluster, which is not conserved in humans. Cluster 7 miRNAs were coordinately downregulated post-chemotherapy and targeted pathways largely overlapping with those regulated by cluster 1 miRNAs (Figure S11).

## Discussion

Our prior work demonstrated that state-transition theory can predict AML development by modeling the distinct yet complementary dynamics of mRNA and miRNA transcriptomes^8^. Here, we extended this framework to investigate how chemotherapy perturbs these transcriptomic state-transitions and identified four key observations.

First, chemotherapy induces a temporal desynchronization between mRNA and miRNA transcriptomic dynamics. Second, this behavior can be captured by a 2D multiomic state-transition framework that models deformation of the leukemic potential landscape during treatment. Third, overexpression of miRNAs from the imprinted *Dlk1-Dio3* locus significantly drives the delayed post-chemotherapy miRNA response. Fourth, we identified a transient activation of immune-activating and metabolic-suppressive pathways during treatment response, which subsequently reverses upon relapse. Together, these findings reveal that chemotherapy induces a temporary regulatory disequilibrium within the AML transcriptome before the system restabilizes during relapse. State-transition frameworks can detect these changes and be applied in the clinic for early assessment of treatment response and possible prediction of outcome.

Post-treatment temporal desynchronization between mRNA and miRNA transcriptome trajectories is a novel finding. This transient divergence suggests that the miRNA transcriptome captures distinct temporal regulatory information rather than merely mirroring mRNA transcriptome. This delayed miRNA response may reflect specific regulatory, or stress-response programs activated downstream of the initial transcriptional response to chemotherapy. The greater stability and slower degradation of miRNAs relative to mRNAs^43,44^ likely also contribute to this temporal delay, allowing miRNA signals to persist even as mRNA trajectories shift toward a healthier state. Because they may be less susceptible to the immediate and potentially nonspecific transcriptional perturbations induced by therapy, miRNA dynamics may serve as a more robust biomarker of treatment response. The divergent mRNA and miRNA dynamics support the innovative hypothesis that mRNA-miRNA desynchronization may function as a biomarker for chemotherapy activity. Future studies are needed to determine whether the timing and magnitude of this desynchronization can serve as early indicators of treatment efficacy and depth of response.

To quantitatively capture desynchronized dynamics, we extended the state-transition framework to model both mRNA and miRNA trajectories. While previous state-transition models have typically characterized cellular differentiation states, cell phenotypes, gene regulatory networks or fitness landscapes in therapeutic resistance^2–5,45–48^, our framework leverages longitudinal multiomic profiling of PB to map chemotherapy-induced transcriptomic state-transitions in AML with known PK/PD effects of chemotherapy. Using this framework, we demonstrated that chemotherapy asymmetrically deforms the leukemic potential landscape, exerting a stronger effect on the mRNA transcriptome than on the miRNA transcriptome. These findings suggest that achieving optimal efficacy may require therapeutic strategies that target miRNA regulatory networks combined with standard chemotherapy. Our treatment vector provides a mathematically rigorous method to quantify the magnitude and direction of therapy-induced landscape deformation, pinpointing which transcriptome is most sensitive to a given intervention. This methodology can be leveraged to design and predict synergistic combinations that drive both transcriptomes toward a health state, establishing a foundational framework for prognostic modeling and treatment design in AML and other hematological malignancies.

To gain biological insight into the delayed miRNA response, we identified a cluster of miRNAs located within the *Dlk1-Dio3* imprinted locus on mouse chromosome 12qF1 (human 14q32) that were strongly upregulated post-chemotherapy. Given the essential roles of miRNAs in AML pathogenesis and their emerging potential as therapeutic targets^49^, overexpression of miRNAs in this locus is particularly notable. In hematological malignancies, coordinated overexpression of *Dlk1-Dio3* miRNAs has been reported in acute promyelocytic leukemia and myelodysplastic syndromes where it correlates with poor clinical outcomes^41,50,51^. Notably, *miR300* residing within the *Dlk1-Dio3* locus and in cluster 1 of our analysis, has been associated with leukemia stem cell (LSC) quiescence and TKI-resistance in CML^52,53^. To the best of our knowledge, this is the first evidence linking the *Dlk1-Dio3* locus with inv(16) AML chemotherapy response. Prior studies indicate that elevated expression of *Dlk1-Dio3* miRNAs suppresses the PI3K/mTOR signaling, a central regulator of cellular metabolism and growth, thereby reducing mitochondrial metabolism and ROS production while preserving hematopoietic stem cell function^54,55^. The functional consequences of PI3K/mTOR suppression in AML are complex and highly context-dependent, including both desired anti-leukemic effects and paradoxical pro-survival adaptations in LSCs that can limit therapeutic efficacy. In this context, our findings support the hypothesis that chemotherapy-induced upregulation of *Dlk1-Dio3* miRNAs drives the coordinated suppression of PI3K–mTOR, OXPHOS, and ROS pathways, potentially safeguarding otherwise vulnerable LSC populations during post-treatment stress. Consistent with this hypothesis, longitudinal transcriptomic analysis revealed that key metabolic pathways such as OXPHOS, glycolysis, and fatty acid metabolism, which are linked to AML progression and therapeutic resistance^30–36^, were downregulated during the miRNA-mRNA desynchronization phase but subsequently reactivated at relapse. Together, these findings highlight metabolic plasticity as an important feature of post-chemotherapy AML recurrence, characterized by transient suppression of oxidative metabolism that is restored during leukemic relapse.

While multiomic approaches are increasingly recognized for improving AML risk stratification and identifying therapeutic vulnerabilities^56–59^, most existing studies lack the temporal resolution required to capture dynamic molecular responses during treatment. Emerging longitudinal multiomic studies have begun to address this limitation but remain constrained by limited sampling and small cohort size^56,60^. In contrast, our approach pairs dense temporal sampling with a mathematical framework grounded in state-transition theory, enabling the explicit modeling of chemotherapy-induced transcriptomic dynamics over time.

In summary, this study provides a longitudinal characterization of chemotherapy-induced transcriptomic dynamics in AML. By revealing a transient desynchronization between mRNA and miRNA dynamics, our findings underscore that chemotherapy acts as a multilayer perturbation whose combined effects define the system-level response. Applying this multiomic state-transition framework enables the quantitative modeling of treatment-induced deformation of the leukemic potential landscape while isolating key regulatory and metabolic programs associated with relapse. Looking forward, this approach can be extended to evaluate other leukemia subtypes and diverse therapeutic modalities, including targeted and epigenetic interventions^61,62^.

## Acknowledgments

This work was supported in part by the National Institutes of Health under award numbers R01CA205247 (Y.-H.K., and G.M.), U01CA250046 (R.C.R., Y.-H.K., and G.M.), U01CA293853

(R.C.R., Y.-H.K., and G.M.), T32CA221709 (J.R.A.), and the Gehr Family Center for Leukemia Research. Research reported in this publication included work performed in the Integrated Genomics, Analytical Cytometry, Animal Resource Center, and Biostatistics and Mathematical Oncology shared resources supported by the National Cancer Institute of the National Institutes of Health under award number P30CA033572. The content is solely the responsibility of the authors and does not necessarily represent the official views of the National Institutes of Health.

## Authorship Contributions

Conceptualization: Y.-H.K., R.C.R., and G.M.; Methodology: R.C.R., J.R.A., Z.C., S.B., D.E.F., L.U., and Y.-H.K.; Data curation: D.O., D.E.F., Z.C.; Investigation: all authors.; Formal analysis: J.R.A., Z.C., R.C.R., Y-H.K., L.U., and Y-H.F.; Visualization: Z.C., R.C.R, and J.R.A.; Writing-original draft: J.R.A., Z.C., R.C.R., G.M., and Y.-H.K.; Writing-review and edit: all authors. Supervision: Y.-H.K., R.C.R., and G.M.; Funding acquisition: Y.-H.K., R.C.R., and G.M.

## Data Sharing Statement

Public deposit in process. Accession number will be provided upon assignment.

## Code Availability

Code to perform the data analyses, mathematical modeling simulations, and to generate all figures is available on GitHub: https://github.com/cohmathonc/AML.Chemo.mRNA.miRNA-manuscript.

