## Supplemental Information for "Multiomic State-Transitions Reveal Post-Treatment Transcriptome Desynchronization in Acute Myeloid Leukemia"

Jennifer Rangel Ambriz^1*^, Ziang Chen^1*^, Yu-Hsuan Fu^2^, David E. Frankhouser^1,2^, Denis O’Meally^1^, Lisa Uechi^1^, Lianjun Zhang^2^, Ying-Chieh Chen^2^, Sergio Branciamore^1^, Jihyun Irizarry^2^, Bin Zhang^2^, Guido Marcucci^2+^, Russell C. Rockne^1+^, Ya-Huei Kuo^2+^

**Supplemental Methods**

### **Mouse model**

Conditional *Cbfb::MYH11* (CM) knock-in mice (*Cbfb^+/56M^/Mx1-Cre*; C57BL/6) were used as a model to recapitulate human inv(16) AML^1^, a common subsets of AML characterized by the rearrangement of chromosome 16 at bands p13 and q22, which generates the chimeric fusion gene *CBFB::MYH11*^2,3^. Expression of CM was induced in 6-8-week-old conditional CM knock-in mice (*Cbfb^56M/+^/Mx1-Cre;* C57BL/6*)* by intraperitoneal injections of polyinosinic–polycytidylic acid [poly (I:C)] (InvivoGen, tlrl-picw-250) at 14 mg/kg per dose every other day for 7 doses. Induced CM knock-in mice develops lethal AML with a median survival of approximately 4 months after induction^1,4^.

### **Flow cytometry analysis**

To estimate the leukemic burden of mice, peripheral blood mononuclear cells (PBMCs) were stained with fluorescence-labeled antibodies (CD117 (c-kit)-APC; eBioscience; 2B8) in phosphate-buffered saline (PBS) with 0.5% bovine serum albumin (BSA) for 15 min at 4 °C. BD LSRFortessa™ X-20 cell analyzer was then used to evaluate the fluorescence intensity of cells. The acquired data were analyzed using Flowjo software 10.6.1.

### **RNA extraction and sequencing**

PBMC pellets collected at each time point were distributed into randomized batches for RNA extraction using the AllPrep DNA/RNA Mini Kit (Qiagen, Hilden, Germany), and their quality and quantity were assessed using a BioAnalyser (Agilent, Santa Clara, CA). Samples with a RIN > 4.0 were included for the following analysis. Samples were then randomized to batches for library preparation, ensuring an even distribution of samples from each timepoint across all sequencing runs.

For the mRNA-seq, sequencing libraries were prepared using the KAPA RNA HyperPrep Kit with RiboErase (Kapa Biosystems, Cat. No. KR1351) according to the manufacturer’s protocol. Library quality was assessed using the Agilent Bioanalyzer with the High Sensitivity DNA assay, and concentrations were determined using the Qubit High Sensitivity DNA Assay (Thermo Fisher Scientific). Libraries were sequenced on the NovaSeq 6000 platform with paired-end 101-bp reads, targeting a nominal depth of 40 million reads. To mitigate batch effects, samples were distributed across eight flow cells in a way that each flow cell contained at least one sample from each mouse and each time point. Real-Time Analysis (RTA) v3.4.4 software was used for base calling, and Illumina bcl2fastq v2.20.0.422 was used to convert base call (BCL) files into FASTQ format.

The nf-core RNASeq pipeline version 3.19.0^5^ was employed to process the raw sequencing reads from the chemotherapy-treated mice, and the untreated CM mice from our previous work^6^. In summary, trimmed reads underwent mapping using Spliced Transcripts Alignment to a Reference (STAR)^7^ against the GRCm38 reference, amended with the human MYH11 fusion transgene. Each library underwent thorough quality assessments, including estimates of library complexity, gene body coverage, duplication rates, and other metrics outlined in the pipeline repository^5^. Abundance was estimated across genomic features using Salmon^8^ and consolidated into a matrix of counts per gene for each mouse at each timepoint. The mRNA-seq dataset will be available soon.

**miRNA-seq library preparation and sequencing**

Library and sequencing methods are as described in Frankhouser et al^9^, and are as follows: All libraries were prepared by the Illumina TruSeq Small RNA protocol with minor modifications following the manufacturer’s instructions. Briefly, for each sample, 280 ng of total RNA was ligated to the sRNA 3′ adaptor (5’-TCTGGAATTCTCGGGTGCCAAGGAACTCC-3’) with T4 RNA Ligase 2, truncated (New England BioLabs) for 1 h at 22°C, and subsequently ligated to a 5′ adaptor: 5’-GUUCAGAGUUCUACAGUCCGACGAUCNNN-3’) with T4 RNA ligase 1 (New England BioLabs) for 1 h at 20°C. The constructed small RNA library was first reverse-transcribed using GX1 (5′- GGAGTTCCTTGGCACCCGAGA) as the RT primer then subjected to PCR amplification for 13 cycles, using the primers GX1 (5′-CAAGCAGAAGACGGCATACGAGAT[NNNNNN]GTGACTGGAGTTCCTTGGCACCCGA GAATTCCA-3’) and GX2 (5′-AATGATACGGCGACCACCGAGATCTACAC[NNNNNNNN]CGACAGGTTCAGAGTTCT ACAGTCCGA-3’), followed by 6% TBE PAGE gel purification with size selection (for targeted small RNAs of 17–35 nt). Individual libraries were prepared using a unique index primer (NNNNNNNN in the GX1 and GX2 primer) in order to allow for pooling of multiple samples prior to sequencing. The purified libraries were quantified using qPCR. Sequencing of paired-end reads was performed on a HiSeq 2500 (Illumina Inc., San Diego, CA), and image processing and base calling were conducted using Illumina's RTA pipeline.

Raw sequencing reads from the current work, and the reads from the untreated CM mice in our previous work^9^, were processed with the nf-core smRNASeq pipeline version 2.3.1^5,10^ using the GRCm38 genome reference (with the parameter --genome GRCm38) and adapters for the Illumina small RNA protocol (by setting --protocol illumina). Briefly, trimmed reads were mapped using bowtie^11^ to miRbase^12^ mature miRNAs (using the parameters -k 50 --best -- strata), and the number of reads mapping to each was counted using samtools stats^13^. Each library was also subjected to extensive quality assessment, including estimation of library complexity, contamination, sequence quality, read length, and depth, among other metrics detailed in the pipeline repository. Mapped reads were merged into a matrix of counts per gene for each sample at each time point and normalized to counts per million (CPM) reads mapped, as implemented in edgeR^14^. The miRNA-seq dataset will be available soon.

### **State-space projection of chemotherapy-treated CM mice into reference state-spaces**

To capture leukemic progression and response to chemotherapy, we used an approach similar to principal component analysis (PCA) to project the transcriptome states of chemotherapy-treated CM mice into previously defined mRNA- or miRNA-based reference state-spaces constructed using untreated CM and control mice^6,9^. To construct the reference state-spaces, we performed PCA via singular value decomposition (SVD) on the mean-centered expression data matrix $\hat{X}=X-\bar{X}$, where $X$ is the log transformed count matrix consisting of all time-series mRNA or miRNA expression data and $\bar{X}$ represents the column mean. The SVD has been extensively used to decompose genomic data^6,9,15–17^ and is given by $\hat{X}=U\Sigma V^{*}$ where * denotes the conjugate transpose. The columns of the unitary matrix $U$ form an orthonormal basis for the sample space and $\Sigma$ is a diagonal matrix containing the singular values. Each column of $V^{*}$ corresponds to specific mRNAs or miRNAs and represents their contribution to different principal components (PCs). Each element in the $V^{*}$ matrix is referred to as the loading value, which quantifies the contribution of an individual mRNA or miRNA to a given PC. Because the PCs capture the major sources of variation associated with leukemia, the loading values indicate the relative contribution of each mRNA or miRNA to disease progression. These columns form orthogonal (independent) directions in a new feature space and are organized by how much variance or information each PC captures in the data. The reference state-spaces were constructed by using the PCs that most strongly correlated with leukemia blast marker *Kit* expression and resulted in the greatest separation between CM and control samples as previously reported^6,9^.

To project transcriptome states from treated CM mice into the reference state-spaces, the mean value $\bar{X}$ and eigengenes $V^{*}$ from untreated CM-control mice were used. First, the gene or miRNA expression data matrix ($X_{Rx}$) of treated CM mice was mean centered to the reference state-space given by $\hat{X}_{Rx}=X_{Rx}-\bar{X}$. Then, the treated CM data were mapped to the corresponding state-space as ${PC}_{Rx}=\hat{X}_{Rx}V$. The first (PC1) and second (PC2) principal components of ${PC}_{Rx}$ define the miRNA- and mRNA-based state-spaces, respectively. These selected PCs correspond to reference state-spaces constructed by untreated CM mice and control mice^6,9^.

#### **Multiomic state-space construction**

To study the mRNA-miRNA dynamics in response to chemotherapy, both data modalities were integrated into a 2D multiomic state-space. Specifically, the mRNA and miRNA transcriptome data obtained from longitudinal PBMC samples were paired based on post-treatment time-points and the respective mRNA and miRNA state-space values were plotted (Figure 2A). The post-treatment time-points are defined from the start of chemotherapy (time = 0 weeks) through 10 weeks post-treatment (time = 10 weeks). To identify any treatment-induced alterations to the mRNA-miRNA relationship relative to the disease state, the mRNA and miRNA state-space values from untreated CM leukemic mice obtained from our previously published work^6,9^ were also included.

#### **Two-dimensional (2D) state-transition model**

To simultaneously model the mRNA-miRNA relationship and identify any treatment-induced perturbations in this relationship, we extended our state-transition framework to a 2D multiomic state-transition model. We modeled the mRNA-miRNA state-transition dynamics as a particle undergoing Brownian motion in a multiomic potential energy landscape with a Langevin equation of the form:

$d\vec{Z}_{t}=-\nabla U_{p}\left( \vec{Z} \right)+\sqrt{2\beta^{-1}}d\vec{B_{t}}.$	(1)

The position of the particle in the potential energy landscape $U_{p}(\vec{Z})$ is defined by the vector $\vec{Z}=\left( z_{1},z_{2} \right)$, where the vector components $z_{1}$ and $z_{2}$ represent the position of the mRNA and miRNA transcriptomes in the state-space, respectively. In component form we have:

$\left\{ \begin{aligned} \frac{dz_{1}}{dt}=-\frac{\partial U_{p}\left( \vec{Z} \right)}{\partial z_{1}}dt+\sqrt{2\beta_{z_{1}}^{-1}}dB_{t} \\ \frac{dz_{2}}{dt}=-\frac{\partial U_{p}\left( \vec{Z} \right)}{\partial z_{2}}dt+\sqrt{2\beta_{z_{2}}^{-1}}dB_{t} \end{aligned} \right.$.

$B_{t}$ is a Brownian stochastic process that is uncorrelated in time $\left\langle B_{i},B_{j} \right\rangle=\delta_{i,j}$ and $\beta_{z_{1}}$ and $\beta_{z_{2}}$ represent the diffusion coefficients associated with the mRNA and miRNA transcriptome trajectories, respectively. A mean squared displacement (MSD) analysis was performed to estimate both diffusion coefficients (Figure S1). Specifically, the MSD for each mouse transcriptome trajectory (mRNA or miRNA) prior to treatment was calculated and a linear fit was applied to the MSD such that the slope of the linear regression was used to estimate the diffusion coefficient. The average slope of the linear regression for the mRNA or miRNA trajectories was used as $\beta_{z_{1}}^{-1}$ and $\beta_{z_{2}}^{-1}$ in the Langevin equation of motion, respectively.

We described the energy potential landscape ($U_{p})$ without any treatment effects as

$U_{p}\left( \vec{Z} \right)=\left( \vec{Z}-\vec{C_{3}} \right)^{2}\cdot\left( \left( \vec{Z}-\vec{C_{1}} \right)^{2}+\gamma\right)$	(2)

where $\vec{C_{1}}=\left( c_{1}, \tilde{c}_{1} \right)$ and $\vec{C_{3}} =\left( c_{3},\tilde{c}_{3} \right)$ denote the health and disease states in 2D, respectively, with the critical points $c_{1}$ and $c_{3}$ corresponding to the mRNA-based state-space, and $\tilde{c}_{1}$ and $\tilde{c}_{3}$, corresponding to the miRNA-based state-space. As previously reported, these critical points were estimated independently from mRNA-seq and miRNA-seq data of untreated CM mice using K-means clustering^6,9^. Briefly, K-means clustering with K=3 was separately performed on the PC2 and PC1 values of the CM samples to identify the three mRNA ($c_{1}, c_{2}$ and $c_{3}$) and miRNA ($\tilde{c}_{1}, \tilde{c}_{2}$ and $\tilde{c}_{3}$) critical points. In particular, the centroids of the cluster K1 and K3 were used to estimate $c_{1}$, $c_{3}$, $\tilde{c}_{1}$ and $\tilde{c}_{3}$, for mRNA and miRNA, respectively. The unstable critical point $c_{2}$ ($\tilde{c}_{2}$ for miRNA) was estimated by maximizing the Boltzmann ratio between $c_{1}$ and $c_{3}$ ($\tilde{c}_{1}$ and $\tilde{c}_{3}$ for miRNA)^6,9^.

Since we identified $\vec{C_{3}} =\left( 0,0 \right)$ and $\vec{C_{1}} =\left( 1,1 \right)$ the equation for the potential landscape becomes

$$U_{p}\left( \vec{Z} \right)= \vec{Z}^{2}\cdot\left( \left( \vec{Z}-\vec{C_{1}} \right)^{2}+\gamma\right).$$

Here, $\gamma$ represents the effect of *Cbfb::MYH11* oncogene activation, introducing an asymmetry in the potential landscape by placing a greater weight on the disease state ($\vec{C_{3}}$), reflecting the higher likelihood of disease progression under untreated conditions (Figure 2B). In the absence of CM, $\gamma= 0$, resulting in a symmetric double-well potential landscape with two local minima and a saddle point in between. Using the second derivative test for functions of two variables we identified the local minima at (0,0) and (1,1) and the saddle point at ($\frac{1}{2},\frac{1}{2}$). To maintain a double-well structure of the potential landscape, the parameter $\gamma$ must be within a range of [0, 0.25). The upper bound is identified from the analytical condition that three critical points are required to maintain the double-well landscape. At $\gamma= 0.25$, the saddle point and one local minimum merge, while for larger values of $\gamma$, the landscape has a one-well structure. This range was further refined to identify the value of $\gamma$ by requiring that trajectories initialized with a position between $c_{2}$ and $c_{3}$ of mRNA state space (also$,\tilde{c}_{2}$ and $\tilde{c}_{3}$ for miRNA), consistent with the data (Figure 1 D,E), relapsed toward $c_{3}$ and $\tilde{c}_{3}$ 10 weeks post-treatment under a single-dose treatment and within 20 weeks under double-dose treatment (Figure S2). This filtering procedure identified a $\gamma$ range from [0.07, 0.12]. From this range, $\gamma=0.08$ was chosen as the value whose simulated trajectories most closely reproduced the qualitative pre-treatment behavior inferred from the data trajectories, which begin between $c_{2}$ and $c_{3}$ ($\tilde{c}_{2}$ and $\tilde{c}_{3}$ for miRNA) and progress toward the $c_{3}$ ($\tilde{c}_{3}$ for miRNA) disease state within approximately 3 weeks (Figure 1, Figure S3).

To account for the effects of treatment, we model the deformation of the multiomic potential field due to chemotherapy by a composition of tensor operators as follows:

$(\phi\circ\psi\circ U_{p})(\vec{Z})=\rho(\vec{Z}\psi\vec{Z}^{T}+E(t))\cdot\left( \left( \vec{Z}-\vec{C_{1}} \right)\phi\left( \vec{Z}-\vec{C_{1}} \right)^{T}+\gamma\right)$	(3)

$= \rho\left( \left( \vec{Z}\psi\vec{Z}^{T} \right)\cdot\left( \left( \vec{Z}-\vec{C_{1}} \right)\phi\left( \vec{Z}-\vec{C_{1}} \right)^{T} \right)+ \gamma\left( \vec{Z}\psi\vec{Z}^{T} \right)+ E(t))\cdot\left( \left( \vec{Z}-\vec{C_{1}} \right)\phi\left( \vec{Z}-\vec{C_{1}} \right)^{T} \right)+ \gamma E(t) \right)$ (4)

Where
 $\left( \psi\circ U_{p} \right)\left( \vec{Z} \right)=\rho\left( \vec{Z}\psi\vec{Z}^{T}+E\left( t \right) \right)\cdot\left( \left( \vec{Z}-\vec{C_{1}} \right)^{2}+\gamma\right)$, $(\phi\circ U_{p})(\vec{Z})=\vec{Z}^{2}\cdot\left( \left( \vec{Z}-\vec{C_{1}} \right)\phi\left( \vec{Z}-\vec{C_{1}} \right)^{T}+\gamma\right)$,
$T$ is the transpose, $\rho$ is a scaling coefficient and $E$ represents the effects of chemotherapy modeled through pharmacokinetic-pharmacodynamic (PK-PD) framework, where the PK component accounts for the drug dose and plasma half-life concentration and the PD compartment translates this drug concentration into the observed drug effect. A detailed definition of $E(t)$ can be found in the following section.

In this model, $\psi$ and $\phi$ are 2x2 tensor operators that map the effects of disease and chemotherapy, respectively, on the multiomic energy potential, altering the landscape post-treatment and are defined as

$\psi=\left[ \begin{matrix} \psi_{1,1} & \psi_{1,2} \\ \psi_{2,1} & \psi_{2,2} \end{matrix} \right]$ and $\phi=\left[ \begin{matrix} \phi_{1,1} & \phi_{1,2} \\ \phi_{2,1} & \phi_{2,2} \end{matrix} \right]$.

The tensor $\psi$ is coupled with $\gamma$, and modulates how the CM activation acts on the mRNA-miRNA interplay after treatment (see second term of Equation (4)). In parallel, $\phi$ is coupled with $E(t)$ and captures how chemotherapy itself deforms the landscape and alters the mRNA-miRNA state-transition dynamics (see fourth term of Equation (4)). Together, $\gamma$ and $E(t)$ represent the CM and chemotherapy-induced effects, while $\psi$ and $\phi$ determine the magnitude and direction of those effects on the mRNA and miRNA dimensions.

The elements $\psi_{i,j}$ and $\phi_{i,j}$ ($i,j=1,2)$ of these tensors increase over time to reach a structure of maximum deformation of the landscape at maximum treatment effect denoted by

$\psi_{maxE}=\left[ \begin{matrix} \psi_{{1,1}_{maxE}} & \psi_{{1,2}_{maxE}} \\ {\psi_{2,1}}_{maxE} & \psi_{{2,2}_{maxE}} \end{matrix} \right]= \left[ \begin{matrix} 1 & 1 \\ 0 & 10 \end{matrix} \right]$ and $\phi_{maxE}=\left[ \begin{matrix} {\phi_{1,1}}_{maxE} & \phi_{{1,2}_{maxE}} \\ \phi_{{2,1}_{maxE}} & \phi_{{2,2}_{maxE}} \end{matrix} \right]= \left[ \begin{matrix} 5 & 0.1 \\ 0 & 3 \end{matrix} \right]$.

The values of these tensor elements were selected to ensure the resulting deformation of the leukemic potential landscape aligned with the direction of the experimental multiomic dynamics (Figure 2C), and that the corresponding single-modality trajectories (Figure 2C) recapitulated the observed response-relapse behavior in both mRNA and miRNA transcriptome trajectories (Figure 1 D,F).

After reaching maximum effect and as the effect of chemotherapy wears off, the tensor elements relax back to a structure that represents untreated conditions:

$\psi_{untreated}=\left[ \begin{matrix} {\psi_{1,1}}_{untreated} & \psi_{{1,2}_{untreated}} \\ \psi_{{2,1}_{untreated}} & {\psi_{2,2}}_{untreated} \end{matrix} \right]= \left[ \begin{matrix} 1 & 0 \\ 0 & 1 \end{matrix} \right]$,

$\phi_{untreated}=\left[ \begin{matrix} {\phi_{1,1}}_{untreated} & \phi_{{1,2}_{untreated}} \\ {\phi_{2,1}}_{untreated} & \phi_{{2,2}_{untreated}} \end{matrix} \right]= \left[ \begin{matrix} 1 & 0 \\ 0 & 1 \end{matrix} \right]$.

To meet these conditions, the tensor elements are defined as follows:

$$\psi_{ij}= {\psi_{ij}}_{untreated}+\left( {\psi_{ij}}_{maxE}-{\psi_{ij}}_{untreated} \right)\left( \frac{E}{\max\left( E \right)} \right),$$

$$\phi_{ij}= {\phi_{ij}}_{untreated}+\left( {\phi_{ij}}_{maxEffect}-{\phi_{ij}}_{untreated} \right)*\left( \frac{E}{\max\left( E \right)} \right).$$

The tensor structure enables separation of modality-specific and interaction effects, with diagonal entries representing direct leukemic or treatment effects on each modality and off-diagonal entries capturing effects on the mRNA-miRNA interplay. Symmetric off-diagonal elements indicate balanced coupling, whereas asymmetric off-diagonal entries imply desynchronized mRNA-miRNA dynamics. In particular, if $\psi_{1,2}>\psi_{2,1}$ or $\phi_{1,2}>\phi_{2,1}$, the leukemic or treatment perturbation acts more strongly on mRNA than miRNA, generating an asymmetric deformation of the potential landscape. As a result, $E(t)$ together with asymmetric tensor entries alter the landscape along the mRNA and miRNA dimensions, shifting the deepest well toward the health state, thereby facilitating transitions from the leukemic state toward the health state once treatment starts (Figure 2C, top panel). The resulting unequal well depths and shifted state-transition critical points suggest distinct responses to treatment in the two data modalities, consistent with the behavior observed in the experimental data (Figure 2C, bottom panel). Therefore, by setting distinct diagonal and non-symmetric off-diagonal entries in $\psi$ and $\phi$, the model simulations recapitulate the altered mRNA-miRNA response dynamics post-chemotherapy that were observed in the data.

Finally, since $\psi$ and $\phi$ are changing over time, the scaling parameter $\rho$ changes accordingly and is defined by:

$$\rho=\rho_{maxE}+\left( \rho_{untreated} - \rho_{maxE} \right)\left( \rho_{untreated} - \frac{E}{max(E)} \right)$$

where $\rho_{maxE}$ and $\rho_{untreated}$ represent the scaling coefficients during maximum drug effect and under untreated conditions, respectively. In this way, $\rho$ will increase after chemotherapy to reach $\rho_{maxE}=0.06$ and as the effects of chemotherapy decay, $\rho$ will decrease towards $\rho_{untreated}=1$.

#### **PK-PD modeling of chemotherapy**

We model the drug effect $E$ at time $t$ by the following Hill equation based on the PK-PD relationship^18,19^:

$$E= \frac{E_{max}\cdot C_{e}}{EC_{50}+C_{e}}.$$

$E_{max}$is the maximum drug effect, $EC_{50}$ is the concentration at which half of the maximum effect is observed and $C_{e}$ is the drug concentration in the effect compartment, which links the PK (dose input) with the PD (dose output) compartment. Specifically, we defined $C_{e}$ by the following differential equation:

$$\frac{dC_{e}}{dt}=k_{e0}(C-C_{e})$$

where $k_{e0}$ is the rate constant for the drug transfer and $C$ is the drug concentration based on the PK model, which takes into account the half-life of cytarabine and daunorubicin ($\lambda_{c}$, $\lambda_{d}$), drug doses ($D_{c}$, $D_{d}$) and the Heaviside step function ($H$) representing the timing of chemotherapy. We defined the half-life for cytarabine and daunorubicin to be $\lambda_{c}=log(2)/0.2404762$and $\lambda_{d}=\log\left( 2 \right)/0.1875$weeks according to literature^20,21^. The drug dose parameters were determined based on the experimentally administered dose multiplied by a mouse average weight of 0.0257 kg such that $D_{c}=1.285$ mg/day and $D_{d}=0.0386$ mg/day. The dynamical model of the “5+3” chemotherapy is given as,

$C=\sum_{i=1}^{5} D_{c}\exp\left( -\lambda_{c}\left( t-t_{i} \right) \right)H\left( t-t_{i} \right)+\sum_{j=1}^{3} D_{d}\exp\left( -\lambda_{d}\left( t-t_{j} \right) \right)H\left( t-t_{j} \right)$.

#### **Defining the treatment vector**

To quantify how chemotherapy perturbs the mRNA-miRNA dynamics and deforms the potential landscape, we compared the untreated $({-\nabla U}_{p}$) and the treatment-perturbed ($-\nabla U_{p}^{Rx}$, where $U_{p}^{Rx}= (\phi\circ\psi\circ U_{p})(\vec{Z})$ at maximum treatment effect i.e. $E=E_{max}$) gradients of the potential fields. We defined a treatment vector $\vec{R}$ as the difference between these two gradients at the spatial location $\vec{Z}^{*}=\left( z_{1}^{*}, z_{2}^{*} \right)$, where the change in direction of force between the two fields is the largest. Mathematically, the largest change in direction between two vector fields corresponds to the minimum dot product, where the dot product is defined as

$$-\nabla U_{p}\cdot-\nabla U_{p}^{Rx}=\left\| -\nabla U_{p} \right\|\left\| -\nabla U_{p}^{Rx} \right\|cos\theta$$

such that $\cos\theta= \left( \frac{-\nabla U_{p}\cdot-\nabla U_{p}^{Rx}}{\left\| -\nabla U_{p} \right\|\left\| -\nabla U_{p}^{Rx} \right\|} \right)$. Since $\cos\theta\in[-1,1]$ is monotonically decreasing on the interval $[0, \pi$], we identified the largest change in direction at $\vec{Z}^{*}$ by identifying the maximum angular difference between two vector fields as follows:

$$\theta_{max}=arccos\left( min\left( \frac{-\nabla U_{p}\cdot-\nabla U_{p}^{Rx}}{\left\| -\nabla U_{p} \right\|\left\| -\nabla U_{p}^{Rx} \right\|} \right) \right) .$$

Values of $\theta_{max}=0^{\circ}$ and $\theta_{max}={90}^{\circ}$ correspond to aligned and orthogonal vectors, respectively, whereas $\theta_{max}={180}^{\circ}$ indicates that the vector fields point in opposite directions at $\vec{Z}^{*}$, representing the maximal difference in the direction of force (Figure S4A). We identified $\theta_{max}={180}^{\circ}$ at $\vec{Z}^{*}=(0.55, 0.18)$ and observed that the vector fields at $\vec{Z}^{*}$ point in opposite directions (Figure S4 B,C).

Finally, the treatment vector at $\vec{Z}^{*}$ was defined as the difference between the untreated and treatment-perturbed normalized vector fields:

$$\vec{R}= \kappa\left( \hat{U}_{p}^{Rx}\left( \vec{Z}^{*} \right)-\hat{U}_{p}\left( \vec{Z}^{*} \right) \right),$$

where $\hat{U}_{p}^{Rx}= \frac{-\nabla U_{p}^{Rx}}{\left\| -\nabla U_{p}^{Rx} \right\|}$ and $\hat{U}_{p}= \frac{-\nabla U_{p}}{\left\| -\nabla U_{p} \right\|}$. The difference field defined over the entire state-space (gray arrows in Figure 2F), and the treatment vector $\vec{R}$ (red vector in Figure 2F), are uniformly scaled by a factor $\kappa=0.1$ for visual clarity. This scaling ensures the vectors are within the state-space bounds between the critical points $c_{3}$ and $c_{1}$ (and $\tilde{c}_{3}$ and $\tilde{c}_{1}$ for miRNA), while preserving directional information. At $\vec{Z}^{*}$, this results in $\vec{R}=(0.19, -0.05)$ with magnitude $\left\| \vec{R} \right\|=0.2$. This treatment vector allows us to quantify how chemotherapy affects the mRNA ($z_{1}$) and miRNA $(z_{2}$) components and deforms the potential landscape.

#### **State-space based sample grouping of mRNA transcriptome trajectories**

Construction of the state-space enabled visualization of leukemia disease progression for individual mice. Because disease progression rates varied across mice, samples from different mice at each timepoint were categorized into three groups based on their spatial proximity to the critical points in the state-space (Figure S5A): pre-treatment (Pre-Rx), maximum therapeutic response (Recovery), and relapse (Relapse). Samples were classified as Pre-Rx upon initial detection of circulating leukemia blasts (cKit+ > 20%), at which point chemotherapy was initiated and samples were closest to the $c_{3}$ leukemic state. State-space trajectories were examined per-mouse to identify the point of maximal treatment response, located closest to $c_{1}$, with the corresponding sample classified as Recovery. Following remission, the sample exhibiting the lowest $c_{3}$ disease state was classified as Relapse.

**miRNA Co-expression modules analysis**

To understand the structure of miRNA disease state space, we used the following approach. First, we selected the top 25% of miRNAs with highest variance among all samples. This step was conducted to minimize noise and identify informative miRNAs only, given the noisy nature of miRNAs. Next, we performed a Weighted Gene Co-expression Network Analysis (WGCNA) to identify co-expressed miRNAs with similar expression dynamic during the disease progression and chemotherapy response. Then, the identified co-expressed miRNA modules were projected back into disease state-space to see each module’s contribution to the state-space during the disease progression trajectory and therapy response.

**Supplemental Tables**

**Supplemental Table 1:** DEG result of different groups

**Supplemental Table 2:** miRNA clustering result

**Supplemental Table 3:** mRNA clustering result

**Supplemental Table 4:** chromosome location of each miRNA

**Supplemental Figures and Legends**

**Supplemental Figure 1**

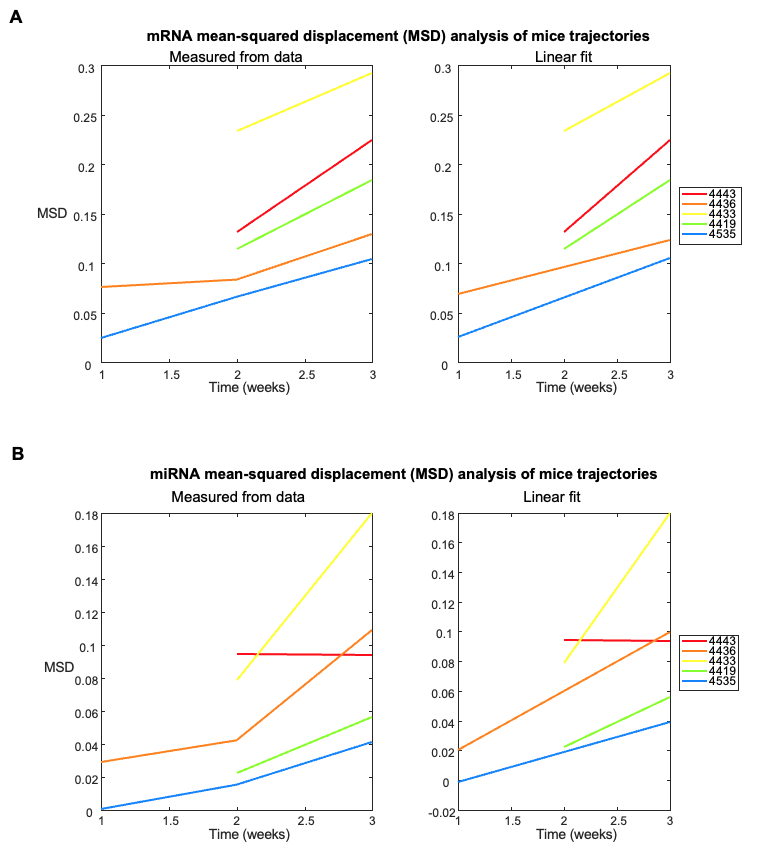

**Figure S1. Mean squared-displacement analysis of mRNA and miRNA transcriptome state-space trajectories.** A mean-squared displacement (MSD) analysis was performed separately on the mRNA and miRNA state-space trajectories to estimate the diffusion coefficients $\beta_{z_{1}}^{-1}$ and $\beta_{z_{2}}^{-1}$ used in the Langevin equation of motion. **A)** The left panel shows the MSD calculated from experimentally derived mRNA trajectories using only pre-treatment time points (from time = -3 weeks to time = 0 weeks). Only mice with available pre-treatment samples were used in this analysis to avoid confounding treatment-induced effects on the dynamics, resulting in n = 5 mice trajectories. Each colored curve corresponds to a single mouse. The right panel shows the corresponding linear fits used to estimate the diffusion coefficient $\beta_{z_{1}}^{-1}$. Assuming Einstein’s relation for diffusive motion, $\beta_{z_{1}}^{-1}$ was estimated as the mean of one-half of the slopes of the linear fits, yielding $\beta_{z_{1}}^{-1}=0.0287 (x^{2}/time),$ which was used in the stochastic simulations. **B)** The same analysis was performed for the miRNA state-space trajectories. Because only positive slopes from the linear fits were used, the slope corresponding to mouse ID 4443 was not included. This yielded an estimated diffusion coefficient of $\beta_{z_{2}}^{-1}=0.0244 (x^{2}/time)$.

**Supplemental Figure 2**

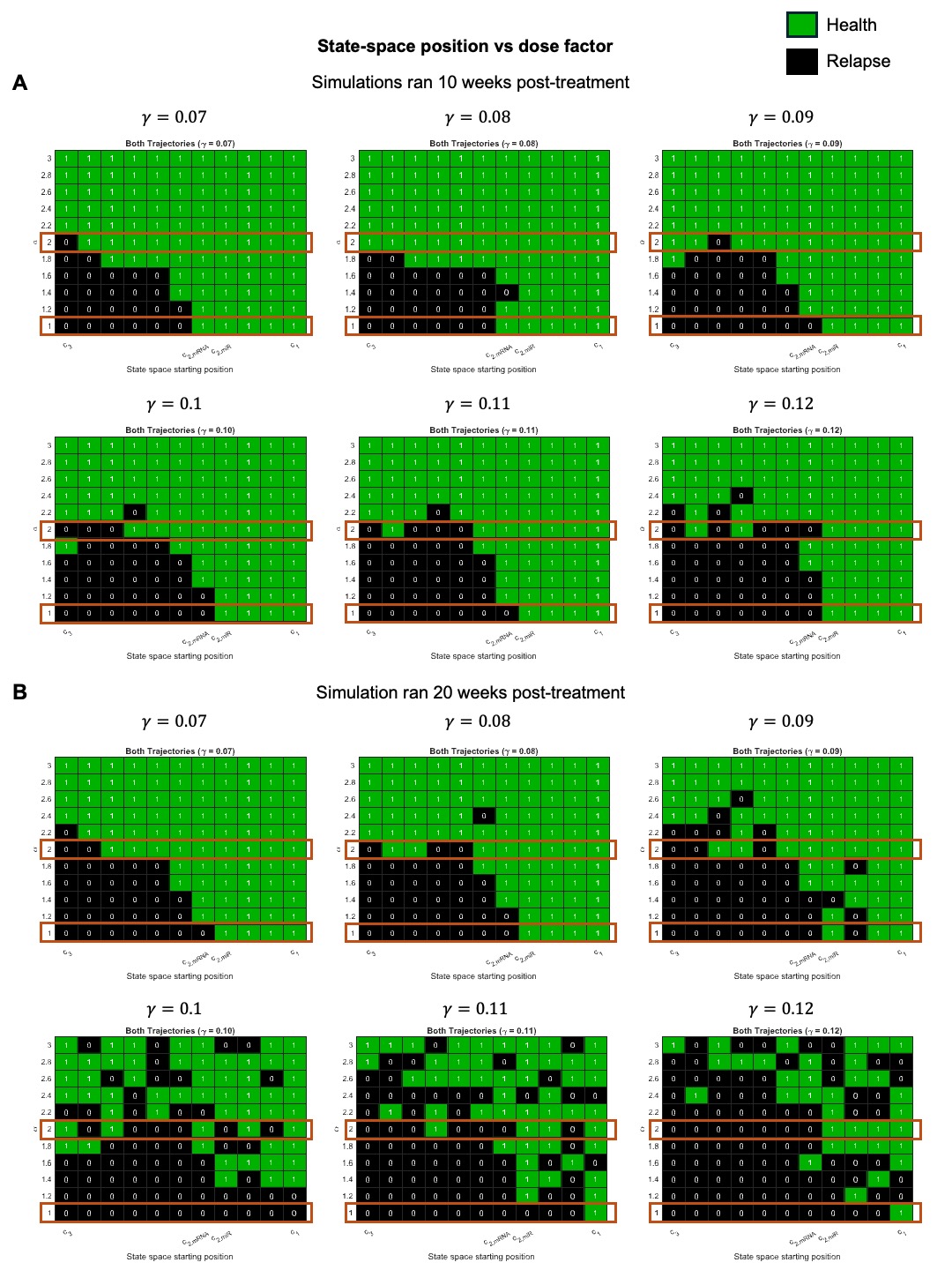

**Figure S2. Analysis to identify a range for parameter** $\boldsymbol{\gamma}$ **based on initial state-space position and dose factor.** Heat maps show the outcomes of the stochastic simulations as a function of changing the state-space starting position $(x_{0})$ and the dose factor ($\alpha$) for different values of $\gamma$. The parameter $\alpha$ scales the drug dose such that $\alpha=2$ corresponds to doubling the dose. Each heatmap is evaluated at a different value of $\gamma$, which represents the leukemic effect in the potential landscape $U_{p}$. The parameter $\gamma$ is constrained to a range of [0, 0.25) to preserve a double-well potential landscape. To further narrow the range of $\gamma$, an analysis was conducted to identify which values of $\gamma$ simulated trajectories relapsed toward $c_{3}$ and $\tilde{c}_{3}$ 10 weeks post-treatment under a single-dose treatment and by 20 weeks under double-dose treatment. **A**) Heatmaps showing the simulation outcomes for simulated trajectories followed for 10 weeks post-chemotherapy. **B)** Heatmaps for simulated trajectories followed for 20 weeks after treatment. Green (value of 1) indicates the simulated trajectory reached the health state ($c_{1}$and $\tilde{c}_{1}$) by the end of the simulation, whereas black (value of 0) indicates relapse toward the disease state ($c_{3}$ and $\tilde{c}_{3}$). Because simulations are stochastic, for each ($x_{0},\alpha$) pair 100 simulations were run and the respective heatmap grid point was colored green if at least 90% of those 100 trajectories reached a health state at the end of the simulation. The single- and double-dose conditions are boxed in orange. These criteria narrowed the range of $\gamma$ to [0.07, 0.12], which is the range shown in panels A and B.

**Supplemental Figure 3**

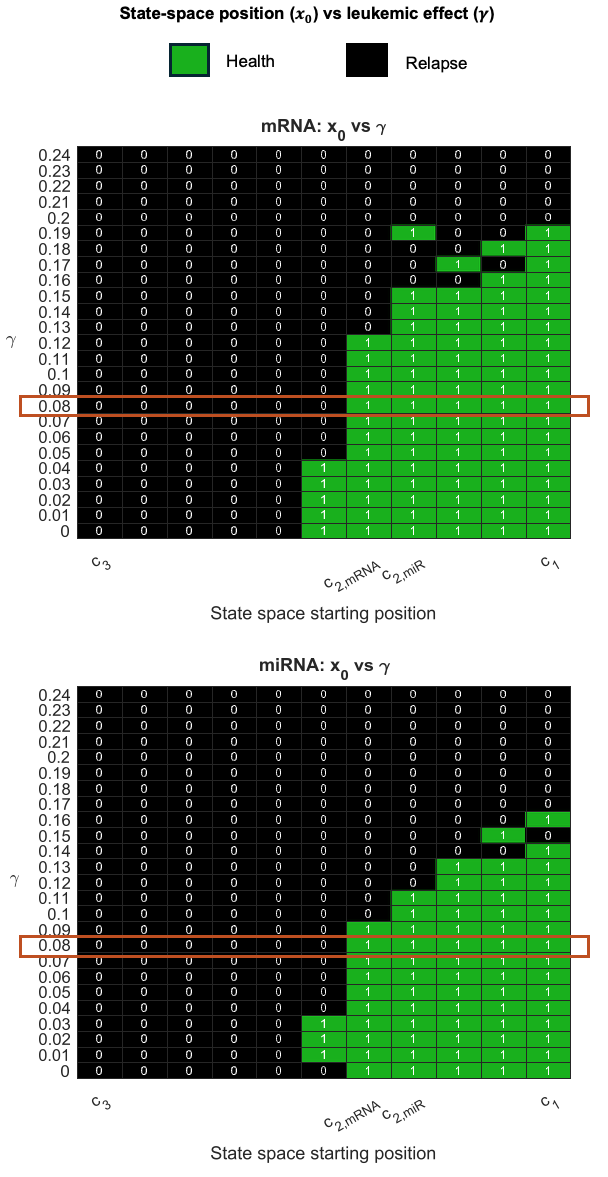

**Figure S3. Analysis of treatment outcome based on initial state-space position and leukemic effect.** Heatmaps show the outcomes of the stochastic simulations for mRNA and miRNA trajectories as a function of changing the state-space starting position $(x_{0})$ and parameter $\gamma$, which represents the leukemic effect in the equation of the potential landscape $U_{p}$. **A)** Heatmap for mRNA trajectories. **B)** Heatmap for miRNA trajectories. The colors green (value of 1) and black (value of 1) represent the treatment outcome at the end of the simulation. Green indicates the simulated trajectory reached the health state ($c_{1}$and $\tilde{c}_{1}$), while black represents disease relapse toward the leukemic state ($c_{3}$ and $\tilde{c}_{3}$). For each ($x_{0},\gamma$) pair, 100 simulations were run and all simulations were followed for 10 weeks post-treatment. A heatmap grid point was colored green if at least 90% of those 100 trajectories reached the health state by the end of the simulation. The results for $\gamma=0.08$ were boxed in orange as this value of $\gamma$ most closely reproduced the qualitative pre-treatment of the data trajectories, which have a state-space starting position between $c_{2}$ and $c_{3}$ ($\tilde{c}_{2}$ and $\tilde{c}_{3}$ for miRNA) and progress toward the $c_{3}$ ($\tilde{c}_{3}$ for miRNA) leukemic state in ~3 weeks.

**Supplemental Figure 4**

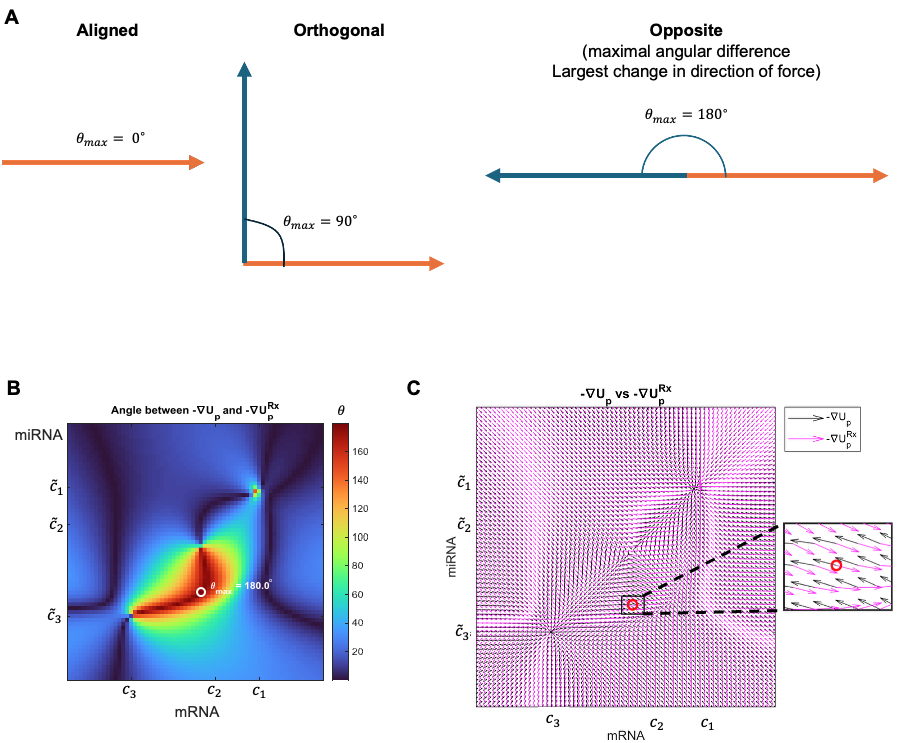

**Figure S4. Computation of the treatment vector using the dot product. A**) A schematic illustrating the geometric interpretation of the dot product between two vectors shown in orange and blue. When two vectors are aligned, the angle between them is $\theta_{max}=0^{\circ}$ and the dot product is the largest, whereas when the vectors are orthogonal, they have zero dot product and $\theta_{max}={90}^{\circ}$. Oppositely oriented vectors have the maximal angular difference of $\theta_{max}={180}^{\circ}$, corresponding to the largest change in direction of force. This geometric intuition motivates the treatment vector analysis, which seeks to identify the spatial location where the change in direction between untreated and treatment-perturbed potential landscapes is the largest, as defined by the maximal angular difference. **B)** Heatmap of the angle $\theta$ between the untreated vector field (${-\nabla U}_{p})$ and the treatment-perturbed vector field during maximum treatment effect ($-\nabla U_{p}^{Rx}$) at every spatial location. The color bar denotes the angular difference in degrees, with a dark red color representing the largest angle ($\theta={180}^{\circ}$) and a dark blue color representing the smallest angle ($\theta=0^{\circ}$). The region circled in white denotes the spatial location $\vec{Z}^{*}=(0.55, 0.18)$ in which the two fields have a maximal angular difference of $\theta_{max}={180}^{\circ}$. **C)** Vector field comparison of ${-\nabla U}_{p}$ (black arrows) and $-\nabla U_{p}^{Rx}$ (magenta arrows). The boxed region is enlarged to show that at $\vec{Z}^{*}=(0.55, 0.18)$, which corresponds to $\theta_{max}={180}^{\circ}$, the vector fields are oppositely oriented and thus, denote the largest change of direction of force and maximal deformation of the landscape due to treatment. The treatment vector $\vec{R}$ is then defined to be the difference between the two fields evaluated at $\vec{Z}^{*}$ (see Methods).

**Supplemental Figure 5**

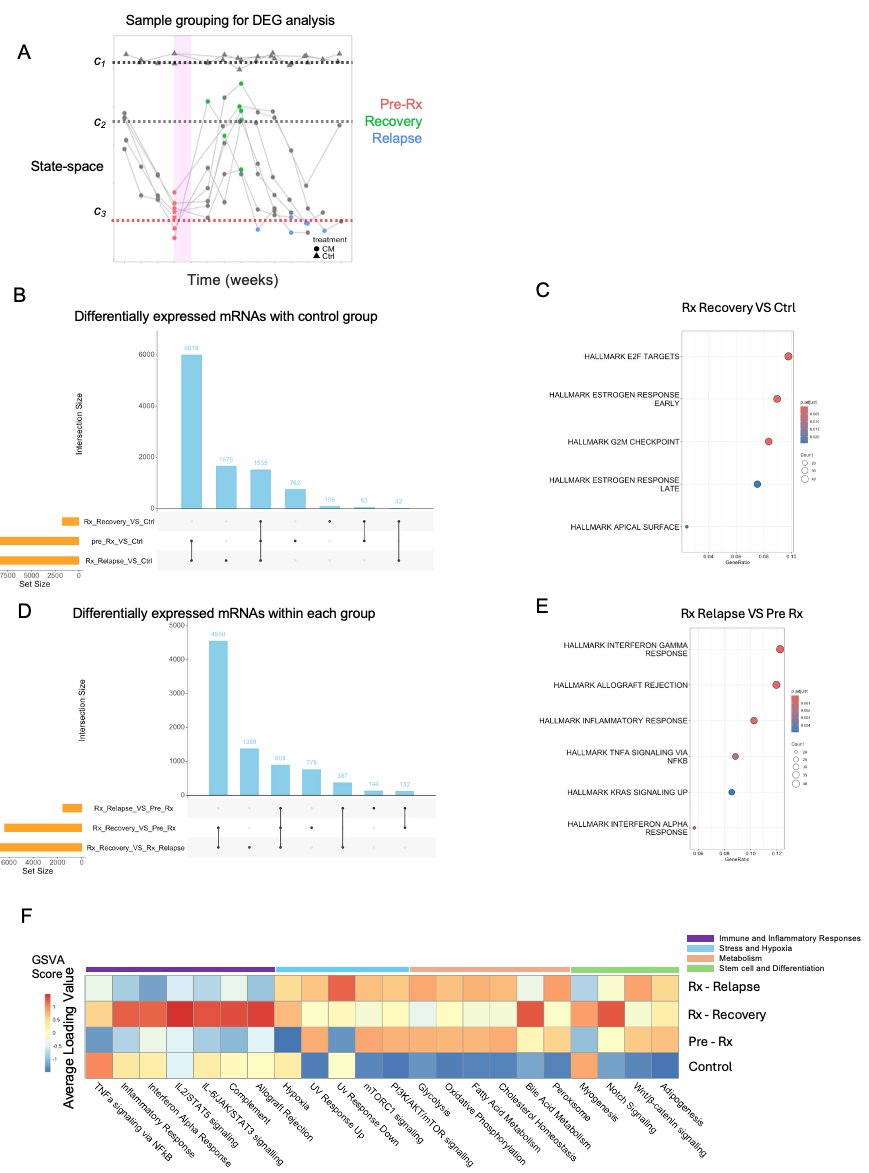

**Figure S5. Differential gene expression (DEG) analysis and Gene Set Variation Analysis (GSVA).** Pathway dynamics at key disease states were analyzed across disease progression using DEG and GSVA. **A)** Samples are plotted in state-space across time (weeks), with the x-axis representing time relative to treatment initiation (week 0) and the y-axis denoting the critical points c₁, c₂, and c₃ with dashed horizontal lines, which define the health, transition and leukemic states, respectively. Each dot represents a CM mouse sample, and each triangle represent a control mice mouse sample. Three disease states were defined based on proximity to the critical points and color-coded by disease state. Pre-Rx (red) represents the pre-chemotherapy leukemic state near c₃, Recovery (green) denotes the maximal treatment response near c₁, and Relapse (blue) represents the return toward c₃. The gray points indicate all other time points with unclassified states. **B)** Horizontal bars indicate the total number of differentially expressed genes (DEGs) identified in each comparison: Rx_Recovery_VS_Ctrl (Recovery), Pre_Rx_VS_Ctrl (Pre-Rx), and Rx_Relapse_VS_Ctrl (Relapse). Vertical bars represent the intersection sizes for each combination of comparisons, as indicated by the connected dots below. The largest intersection (6,018 genes) is unique to the Relapse versus Pre-Rx comparison, reflecting the greatest transcriptional perturbation occurring during the recovery phase following treatment. Smaller intersections of 1,676 and 1,535 genes represent DEGs shared between or unique to other comparisons, while minimal overlaps (762, 106, 63, and 42 genes) are observed across multi-group intersections. **C)** GSEA result of the DEGs in Rx_Recovery_VS_Ctrl **D)** Horizontal bars indicate the total number of differentially expressed genes (DEGs) identified in each comparison: Rx_Relapse_VS_ Pre_Rx , Rx_Recovery_VS_ Pre_Rx, Rx_Recovery_VS Rx_Relapse(. Vertical bars represent the intersection sizes for each combination of comparisons, as indicated by the connected dots below. **E)** GSEA result of the DEGs in Rx_Relapse_VS_ Pre_Rx. **F)** Heatmap displays the average GSVA loading values for hallmark pathways across four disease states (Control, Pre-Rx, Rx-Recovery, and Rx-Relapse) with red indicating positive enrichment and blue indicating negative enrichment. Pathways are grouped into four functional categories annotated along the top: Immune and Inflammatory Responses (purple), Stress and Hypoxia (light blue), Metabolism (yellow), and Stem cell and Differentiation (green).

**Supplemental Figure 6**

**Figure S6. Distribution of normalized state-space values for miRNA (blue) and mRNA (green) across four time points (Week 0, 2, 3, and 4).** Boxes represent the interquartile range with the median indicated by the center line. Statistical significance was assessed using ANOVA test with different groups at difference time points. The result showing a significant difference between mRNA and miRNA within W0 and W2, indicated the existence of desynchronization two weeks post-chemotherapy.

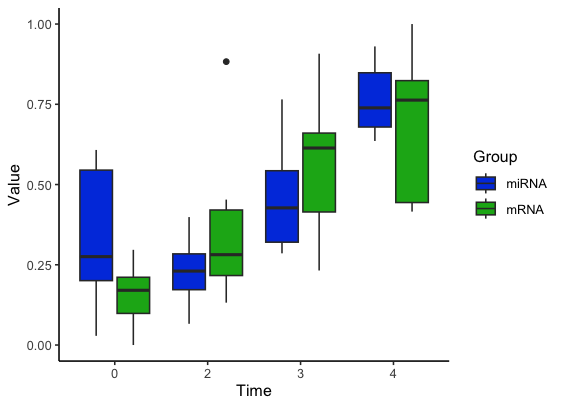

*

ns

ns

Normalized state-space

**Supplemental Figure 7**

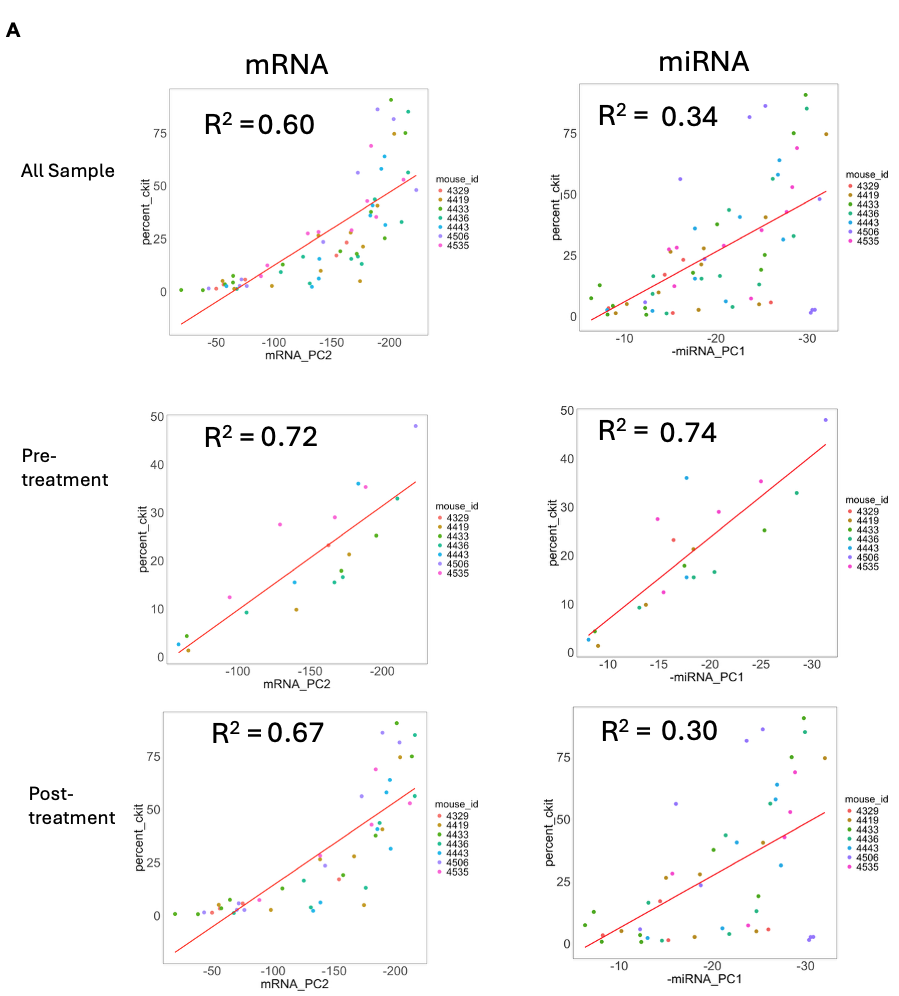

**Figure S7. Correlation of %cKit+ with mRNA and miRNA state-spaces. A)** Percent of cKit+ cells (%cKit+) as determined by flow cytometry plotted against the mRNA (left column; PC2) and miRNA (right column; PC1) state-spaces constructed using principal component analysis (PCA). Each point corresponds to a time point sample colored by mouse ID, with the same color used for all time points from any given mouse. The red lines indicate linear regression fits. When considering all time point samples (top row), the mRNA state-space shows a stronger correlation with %cKit+ (R² = 0.60) than the miRNA state-space (R² = 0.34). When stratified by treatment condition, both state-spaces exhibit stronger correlations pre-treatment (middle row; miRNA R² = 0.74, mRNA R² = 0.72). Post-treatment (bottom row), the correlation decreases for miRNA (R² = 0.30) but remains relatively strong for mRNA (R² = 0.67). Pre-treatment conditions include all time points up to treatment administration (time = 0 weeks), while post-treatment includes all time points after time = 0 through 10 weeks after chemotherapy.

**Supplemental Figure 8**

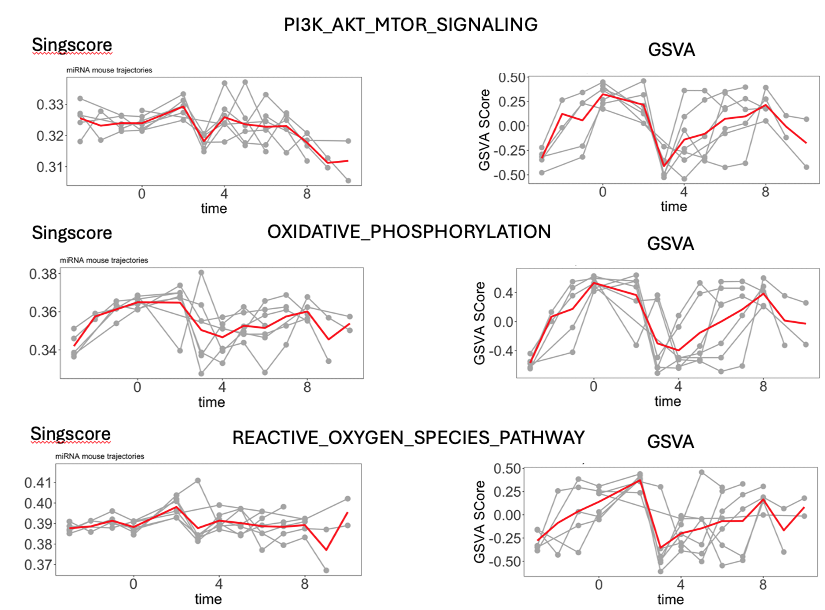

**Figure S8. Temporal dynamics of metabolic pathway activity assessed by Singscore and Gene Set Variation Analysis (GSVA) across disease progression and post-treatment response.** Time-related dynamics of three major metabolic pathways, PI3K/AKT/mTOR signaling, oxidative phosphorylation (OXPHOS), and reactive oxygen species (ROS) pathway, scored by two complementary methods: Singscore, a rank-based method that scores each sample independently, and GSVA, which estimates pathway activity as a continuous enrichment score across the expression matrix. The x-axis represents time in weeks relative to treatment initiation. Gray lines represent individual mouse trajectories, and the red line indicates the mean trend across all samples.

**Supplemental Figure 9**

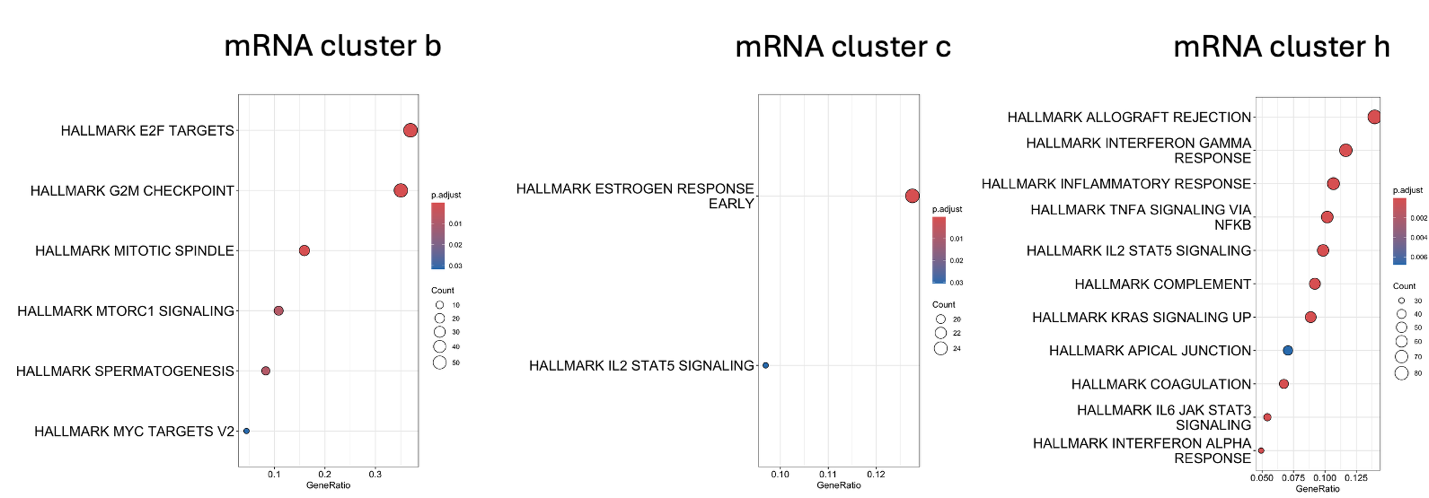

**Figure S9. Pathway enrichment analysis for mRNA clusters b, c, and h.** Hallmark gene set enrichment for mRNA clusters b, c, and h. The gene ratio on the x-axis reflects the proportion of enriched genes per pathway. Dot size corresponds to gene count. Dot color indicates adjusted p-value (p.adjust), with a red color denoting higher significance.

**Supplemental Figure 10**

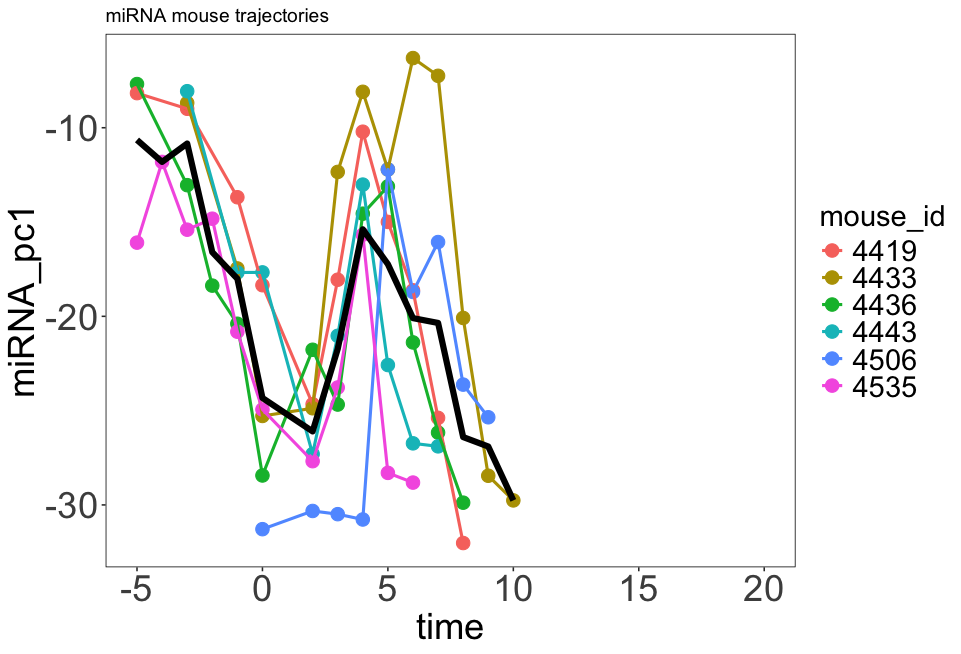

Original miRNA trajectory

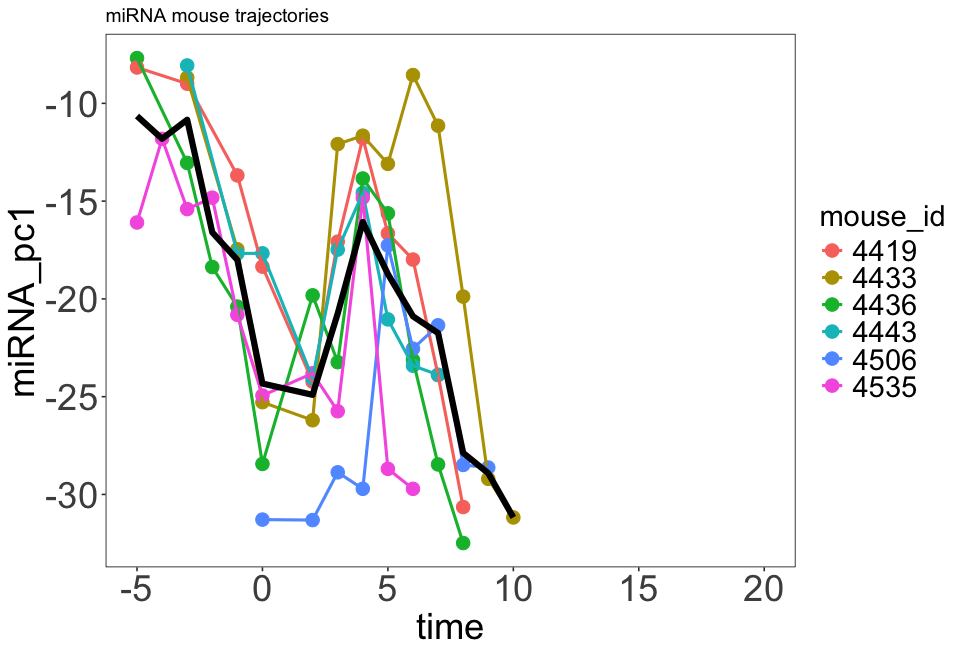

Replace Cluster 1 after Rx with W0

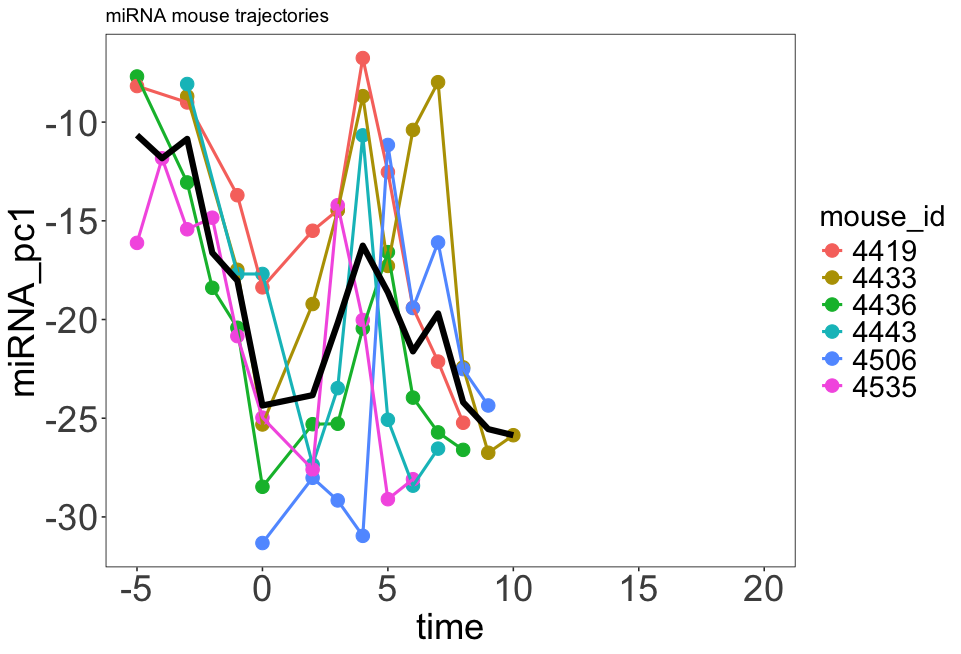

Replace Cluster 7 after Rx with W0

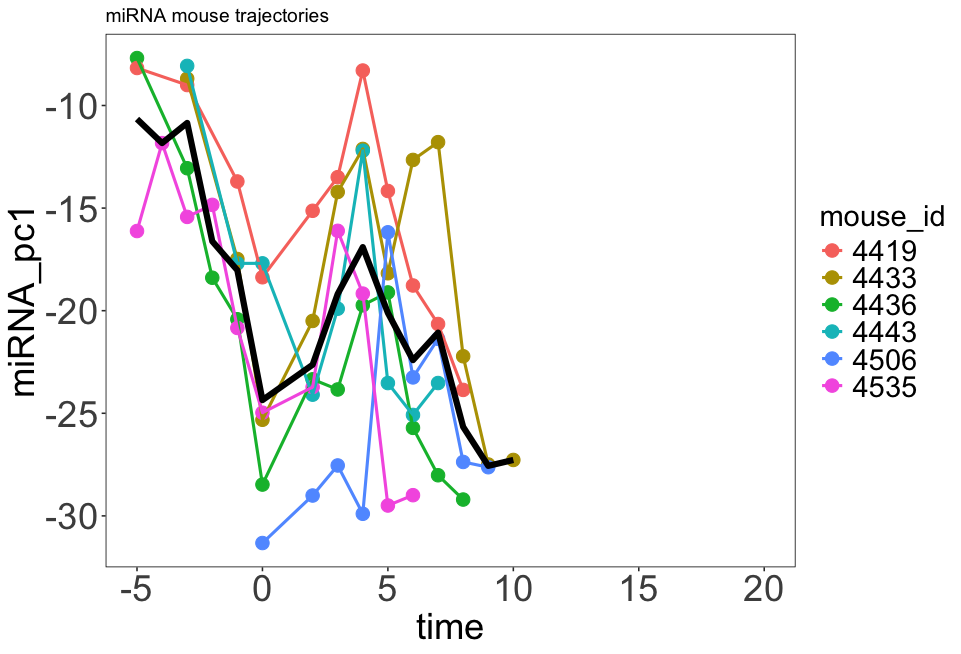

Replace Cluster 1&7 after Rx with W0

**Figure S10. State-space analysis of miRNA trajectory dynamics following chemotherapy with computational removal of miRNA clusters 1 and 7.** Longitudinal miRNA state-space trajectories are shown for individual mice before and after in silico removal of miRNA clusters 1 and 7. In each panel, the black line represents the mean trajectory across mice and summarize the transcriptome dynamics. Chemotherapy treatment was initiated at W0 and denoted in pink. In the original trajectories (top left), all mice exhibit a coordinated decline approximately two weeks after chemotherapy administration at W0 (time = 0), followed by a transient rebound and subsequent relapse, reflecting a conserved dynamical behavior across mice trajectories. Because the delayed response emerges ~2 weeks after treatment, data from mice lacking measurements at W2 (time = 2) and W3 (time = 3) were excluded from the average trajectory (black line) to ensure consistent temporal comparisons across all mice trajectories. Accordingly, mouse ID 4329 was removed. Computationally setting the expression levels of miRNAs in cluster 1 (top right), cluster 7 (bottom left) or both clusters (bottom right) to their respective pre-treatment levels mitigates the delayed response, with the combined removal of both clusters results in the complete disappearance of the delayed effect.

**Supplemental Figure 11**

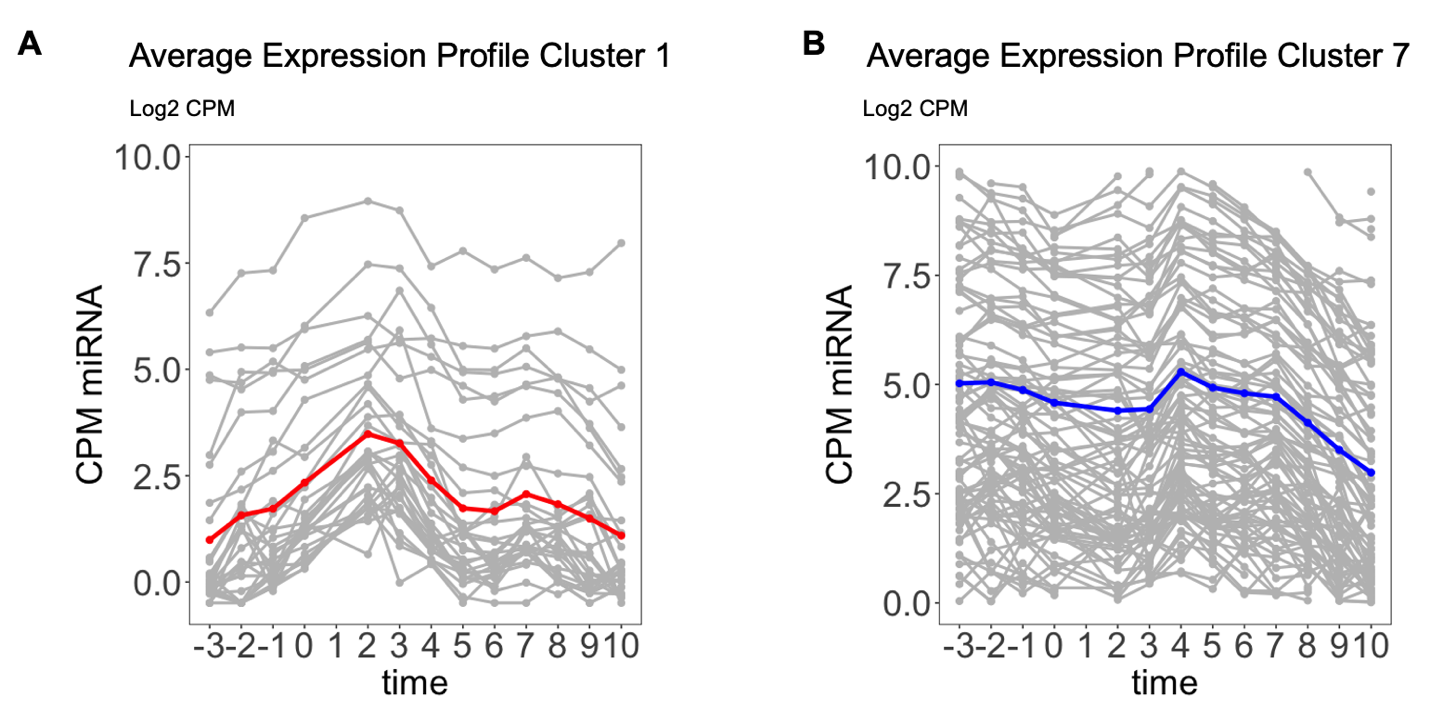

**Figure S11. Average expression profiles of miRNAs in clusters 1 and 7.** Average log2 counts per million (CPM) expression over time for miRNAs in **A)** cluster 1 and **B)** cluster 7. Each gray line represents the expression profile of an individual miRNA within the cluster. The mean expression profile for cluster 1 and 7 are shown in red and blue, respectively. Time (in weeks) is shown relative to treatment, with time = 0 indicating the start of chemotherapy. The average expression of cluster 1 miRNAs displays an increase after chemotherapy (time = 0), peaking around 2-3 weeks post-treatment, followed by a decrease in expression at later time points. In contrast, the average expression of miRNAs in cluster 7 exhibits a modest decrease after chemotherapy, followed by a gradual increase around week 4 and a subsequent decrease in expression in the later time points.

**Supplemental Figure 12**

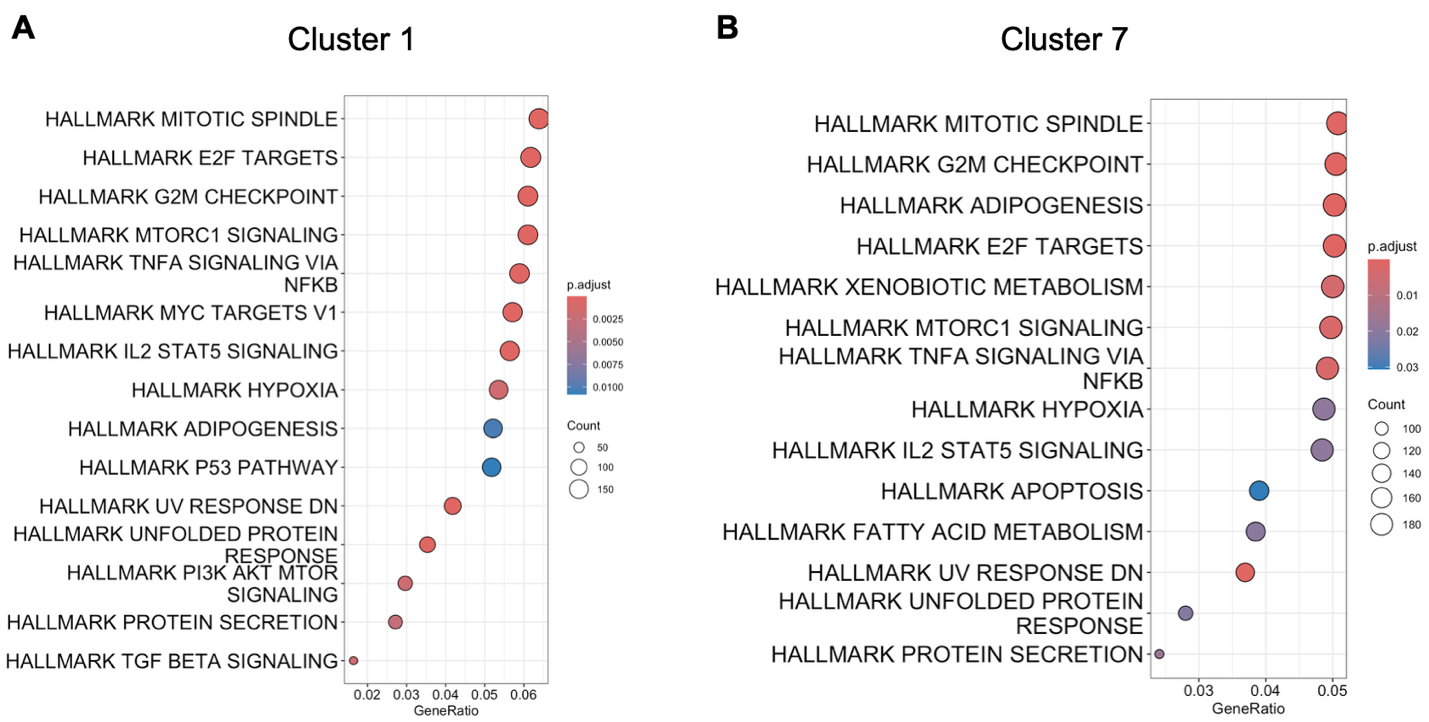

**Figure S12. Pathway enrichment analysis of experimentally validated mRNA targets of miRNAs from clusters 1 and 7.** Experimentally validated mRNA targets of miRNAs from clusters 1 and 7 were identified using miRecords, miRTarBase, and TarBase databases. Gene set enrichment analysis (GSEA) was performed on the gene targets to determine enrichment of Hallmark pathways. Dot plots display significantly enriched Hallmark gene sets for **A)** cluster 1 and **B)** cluster 7. Pathways are ranked by gene ratio (x-axis), which is defined as the proportion of genes in each cluster that overlap with a given pathway. Dot size represents the number of target genes contributing to each pathway, and color indicates the adjusted p-value (p.adjust), with warm colors reflecting higher statistical significance.

**Supplemental Figure 13**

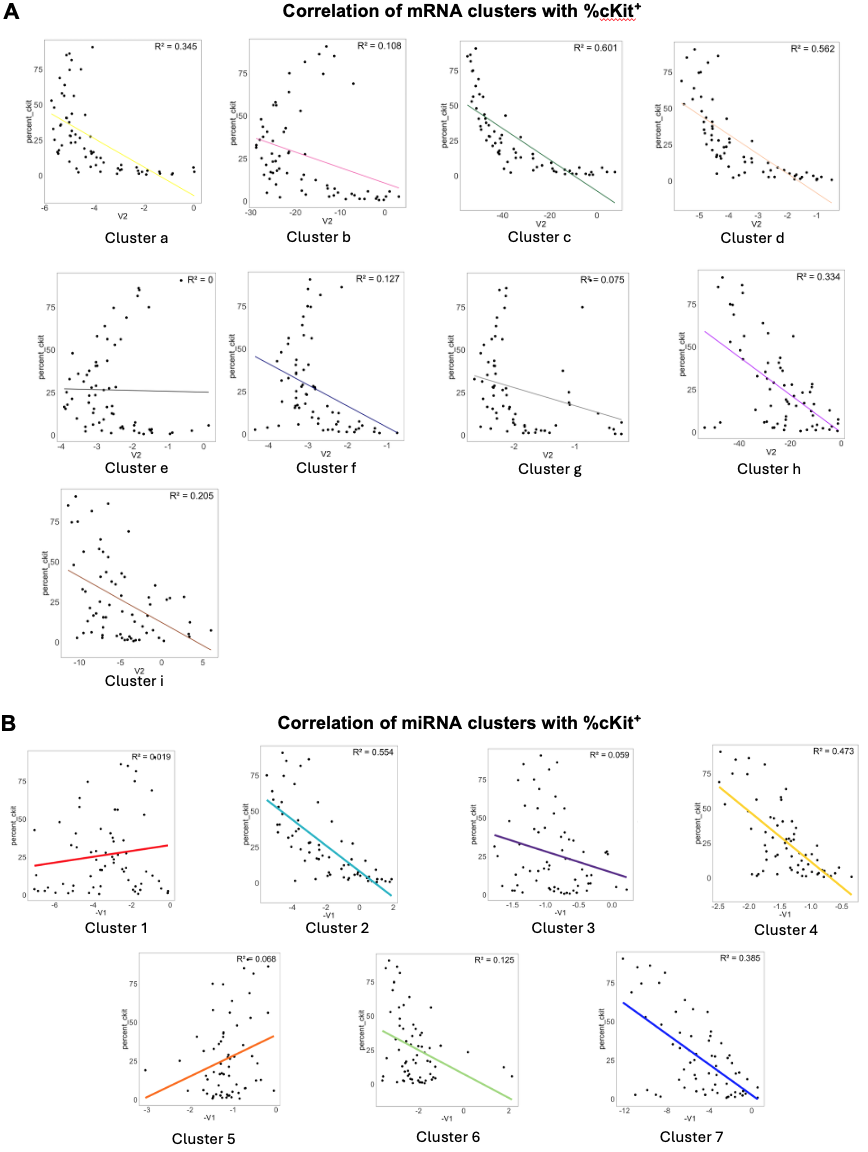

**Figure S13. Correlation between mRNA and miRNA cluster expression with %cKit+ across all samples. A)** The mean expression of each mRNA cluster (x-axis) plotted against the mean cKit+ percentage (y-axis) across all samples. Each panel corresponds to one mRNA cluster (Clusters a-i), with the coefficient of determination (R²) indicated in the upper right corner. Clusters c (R² = 0.598), d (R² = 0.645), h (R² = 0.513) and i (R² = 0.52) demonstrate the strongest positive correlations with %cKit+, while all other clusters display weaker correlations. **B)** The correlation between the mean expression of each miRNA cluster (x-axis) and mean cKit+ percentage (y-axis) across samples. Each panel represents an independent miRNA cluster (Clusters 1–7), with R² value in the upper right corner. Clusters exhibiting negative slopes (Clusters 2, 3, 4, 6, and 7) demonstrate inverse associations between miRNA cluster expression and cKit+ activity. Clusters 2 (R² = 0.717), 4 (R² = 0.738) and 7 (R² = 0.511) demonstrate the strongest correlations with %cKit+, while clusters 1 (R² = 0.131) and 5 (R² = 0.199) display modest positive associations. Regression lines are color-coded per cluster for visual distinction.

**Supplemental Videos**

**Video 1. Two-dimensional (2D) state-transition model simulation.** Left panel: Time evolution of a 2D particle with position $\vec{Z}=\left( z_{1},z_{2} \right)$ in a multiomic state-space. The trajectory represents the simulated mRNA-miRNA dynamics. The initial position of the simulated mRNA-miRNA trajectory is denoted in black, and the final position is denoted in cyan. Contour lines represent the underlying multiomic potential landscape. The landscape deforms over time due to the effects of chemotherapy, resulting in time-dependent changes of the contour lines. Right panel: Corresponding leukemic potential landscape, with projected contour lines shown on the base plane. The potential landscape has two wells to represent the state of disease $(c_{3},\tilde{c}_{3})$ and the state of health $(c_{1},\tilde{c}_{1})$. Upon treatment onset, the landscape is deformed.
